# *Vibrio cholerae* ParE2 toxin modulates its operon transcription by stabilization of an antitoxin DNA ruler

**DOI:** 10.1101/2021.03.22.436508

**Authors:** Gabriela Garcia-Rodriguez, Yana Girardin, Ranjan Kumar Singh, Alexander N. Volkov, Albert Konijnenberg, Frank Sobott, Daniel Charlier, Remy Loris

## Abstract

The *parDE2* operon of *Vibrio cholerae* encodes a type II TA system, which is one of three loci in the superintegron of small chromosome II that show modest similarity to the *parDE* operon of plasmid RK2. ParE2, like plasmid RK2-encoded ParE, inhibits DNA gyrase, an essential topoisomerase that is also the target of quinolone antibacterial agents. Mechanistic understanding on ParE2 toxin inhibition by direct interaction with its cognate antitoxin and transcriptional autoregulation of the TA system are currently lacking. ParD2, the ribbon-helix-helix (RHH) antitoxin, auto-represses the *parDE2* promoter. This repression is enhanced by ParE2, which therefore functions as a transcriptional co-repressor. Here we present protein-DNA interaction studies and high-resolution X-ray structures of the ParD2:ParE2 complex and isolated ParD2 antitoxin, revealing the basis of toxin inhibition and autoregulation of the TA operon by conditional cooperativity. Native mass spectrometry, SAXS and MALS studies confirm the presence of different oligomerization states of ParD2 in solution and the role of the DNA-binding hexameric ParD2_6_:ParE2_2_ assembly in transcriptional repression.

## Introduction

Plasmid-encoded stabilization elements were first discovered when studying mechanisms responsible for controlled maintenance of extrachromosomal elements, including naturally occurring plasmids (for a review see Gerdes et al., 2000). Partition regulatory systems include stable toxins inhibited by their labile counterparts under constant production. In the event of plasmid or gene loss, the long-lived toxin now liberated from its antidote is able to target an essential cell process culminating in death of plasmid (locus)-free daughter cells, a mechanism termed post-segregational killing (Gerdes et al., 1986). The *ccd* locus on the F episome (Jaffé et al., 1985) and the *hok*/*sok* locus on plasmid R1 (Gerdes et al., 1986) were the first two of this type of genetic systems to be discovered, although their exact molecular mechanisms and regulation remained elusive at the time. Subsequently, other plasmid-stabilizing systems were discovered in different mobilomes, including two divergently transcribed operons in the broad host-range plasmid RK2, denoted *parCBA* and *parDE*. The latter was found to be essential for plasmid stabilization (Roberts and Helinski, 1992). The toxic product of the *ccd* and *par* systems (CcdB and ParE), responsible for plasmid-free cell killing, was later found to poison the same cellular target: DNA-topoisomerase II or gyrase (Bernard and Couturier, 1992; Jiang et al., 2002). Nearly all bacterial and archaeal chromosomes sequenced to date contain one or multiple copies of Toxin-Antitoxin (TA) loci (Pandey and Gerdes, 2005; Makarova et al., 2009), with some species, like the persistent pathogen *Mycobacterium tuberculosis*, showing more than 80 predicted TA loci on its chromosome (for a review see Ambre et al., 2014). They often localize within mobile DNA elements, such as integrons platforms within transposons, conjugative plasmids or (defective) prophages. These observations, plus other findings such as the upregulation of some TA systems under stress conditions, led to the proposed role of their involvement in stress response mechanisms (for a review see Gerdes et al., 2005). Consequently, the physiological roles and regulation of these chromosomal loci have been the subject of extensive research, although controversy regarding their true biological functions still exists today (LeRoux et al., 2020; Fraikin et al., 2020).

*Vibrio cholerae*, the Gram-negative bacterium that causes cholera (whose genome, like that of all *vibrios*, is divided between two chromosomes), also displays a high number of TA systems in its second, smaller chromosome. TA operons are clustered as gene cassettes in the superintegron (SI) of chromosome II (Iqbal et al., 2015; Pandey, 2005; Rowe-Magnus et al., 2003), and they stabilize SIs massive arrays in absence of selective pressure (Iqbal et al., 2015; Szekeres et al., 2007).

Typically, gene cassettes on integron platforms are promoterless. Their expression is driven by the position relative to the Pc promoter - the latter being internal to the *intI* gene (encoding integrase) of the integron (Collis and Hall, 1992, 1995). Analysis of the SI on chromosome II of *V. cholerae* led to the identification of 13 putative TA loci (Rowe-Magnus et al., 2003; Pandey and Gerdes, 2005), and later to four more loci localized within the SI plus one HipAB-related operon, which does not localize within a mobile genetic element (Iqbal et al., 2015). Importantly, all of the 18 identified TA loci-including three *parDE*, two *higBA* and the *phd-doc* cassettes that had already been confirmed (Christensen-Dalsgaard and Gerdes, 2006; Guerout et al., 2013; Yuan et al., 2011)-were shown to be functional TA systems in *V. cholerae* (Iqbal et al., 2015). Interestingly, these cassettes seem to contain their own promoter (for a review see Cambray et al., 2010), which strengthened the hypothesis of their role in the stabilization of these long genomic regions. Furthermore, these regions are generally non-essential for bacterial survival, especially under vegetative growth, but rather enhance virulence and confer adaptive potential to environmental pressures (Yuan et al., 2011; Christensen-Dalsgaard and Gerdes, 2006).

Among these 17 identified TA cassettes on *V. cholerae* SI, three are present in two identical copies: two *parDE* loci (*parDE1* and *parDE3*, Yuan et al., 2011), two *relBE* loci (*relBE2* and *relBE4*) and the newly identified VCA0318-VCA0319 and VCA0481-VCA0482 loci (Iqbal et al., 2015), thus rendering 14 different cassettes. Cross-reactivity between all 14 identified TA systems was addressed, including the paralogous sets of *parDE*, *relBE* and the newly identified 318-319/481-482, by co-expression of all possible 196 toxin-antitoxin pairs in *E. coli* and rescue from toxin-exerted toxicity by antitoxin expression. Remarkably, no cross-interaction was observed between all 87 non-cognate pairs (Iqbal et al., 2015).

The *parDE2* operon on the SI of *V. cholerae*, along with paralogous *parDE1-3* loci, shows low similarity to *parDE* from plasmid RK2. *parDE2* encoded toxin, ParE2, like its RK2 ParE homolog, targets DNA gyrase. Interestingly, gyrase is targeted on different sites by ParE2 and CcdB, which suggests that ParE2 poisons gyrase via a different mechanism (Yuan et al., 2010). ParD2 neutralizes ParE2 toxicity through *in vivo* formation of a ParD2:ParE2 protein complex (Yuan et al., 2010). Even though the *parDE* family of TA systems is often found in multiple copies on bacterial chromosomes (Pandey and Gerdes, 2005; Leplae et al., 2011), neither the regulatory mechanism of toxin activity nor the autoregulation mechanism of the TA loci is known to date. ParD antitoxins are highly specific for their cognate toxins. In a study where more than 20 chromosomally encoded ParD:ParE pairs from eight different bacteria where analyzed for cross-interactions, only 3 % of non-co-operonic pairs were found to interact, and these never originated from the same species (Aakre et al., 2015). This demonstrates the high specificity of ParD-ParE pairs, i.e., ParD antitoxins can only neutralize their cognate toxins.

As in many other antitoxins, the N-terminal part of ParD bears the DNA binding domain, which is crucial for transcriptional autoregulation of the operon, while the C-terminus is involved in toxin neutralization by direct binding to ParE (Aakre et al., 2015; Dalton and Crosson, 2010; Oberer et al., 2007). The autoregulatory function of ParD antitoxins binding to their operator region also shows high specificity, since transcription of the paralogous *parDE1-3* and *parDE2* loci from *V. cholerae* SI is independently regulated by their cognate antitoxin (Yuan et al., 2011). Conditional cooperativity (CC) as a transcriptional regulation mechanism emerged as a potentially common feature of TA loci with the study of the *relBE* system (Overgaard et al., 2008, 2009). CC has been studied with the TA operons *vapBC* from *Salmonella enterica* (Winther and Gerdes, 2012), *phd/doc* from plasmid P1 (Garcia-Pino et al., 2010, 2016; Magnuson and Yarmolinsky, 1998) and *ccdA/ccdB* from episome F (Afif et al., 2001; Vandervelde et al., 2017). This mechanism underpins the role of the toxin as modulator of transcriptional levels of the entire operon by acting as a corepressor/derepressor of operator-binding antitoxin upon fluctuations of the toxin:antitoxin (T:A) ratio. Corepression is observed at low T:A ratios, where the binding of the toxin increases the avidity of the antitoxin for the operator DNA. Derepression occurs when T:A ratios increase, typically above 1 and a transition occurs to a low DNA-affinity antitoxin state, or by switching to a high-affinity T:A interaction that results in a complex with a different architecture, which is unable to repress the operon (Overgaard et al., 2008; Garcia-Pino et al., 2010; Garcia-Pino et al., 2016).

Here we describe the X-ray structures of *V. cholerae* ParD2 antitoxin and ParD2:ParE2 complex, unveiling the basis for ParD2 DNA and toxin binding. We show that ParD2 has the ability to form higher-order oligomers and its intrinsically disordered C-terminus folds upon binding ParE2 toxin. Native mass spectrometry and SAXS studies confirmed the unusual stoichiometry of the ParD2:ParE2 complex in solution. Computational modeling combined with X-ray crystallography and SAXS confirms the intrinsic disorder of the C-terminus of ParD2. Furthermore, protein-DNA complex formation was studied by Electrophoretic Mobility Shift Assays (EMSAs), the binding site for ParD2 in the *V. cholerae parDE2* promoter region delimited by enzymatic footprinting, and base and groove specific interactions important for complex formation identified by chemical premodification binding interference and methylation protection experiments. EMSAs and other biochemical assays indicate the presence of conditional cooperativity in the transcriptional regulation of this system. The high-resolution structures of isolated ParD2 and ParD2:ParE2 complex reveal the basis for conditional cooperativity and the different stoichiometric states populated by the TA complex and antitoxin.

## Results

### ParD2:ParE2 complex exists as a 6:2 hetero-octamer in solution

The stoichiometry of the ParD2:ParE2 complex was first assessed by analytical gel filtration and native mass spectrometry. The theoretical masses for the monomers of ParD2 and ParE2 are 8,9 and 12,4 kDa, respectively (including the C-terminal 6xHis tag of ParE2). However, the estimated molecular mass by gel filtration of 70 kDa does not correspond to the theoretical mass of a heterodimer (21,3 kDa) nor to that of a heterotetramer. Therefore, native mass spectrometry experiments were performed to determine the stoichiometry of the complex in solution. In a first step, the purity of the preparation was assessed by measuring a spectrum under denaturing conditions, where all detected peaks corresponded to the exact masses of the full length and intact ParD2 and ParE2 protomers. Subsequently, the complex was measured under “native” conditions, which relies on the gentle transfer of folded proteins from a volatile aqueous buffer (ammonium acetate) into the vacuum of the spectrometer using nano-electrospray ionization (nano-ESI). The peaks with the highest intensity in the spectrum were detected at the mass-to-charge (m/z) ratios corresponding to the exact mass of the 6:2 association between ParD2 and ParE2 (77,85531 kDa). Peaks corresponding to the association of two of these 6:2 complexes were also detected (156,47705 kDa), as well as peaks of ParD2 monomer and dimer. This stoichiometry is surprising, since it has not been previously reported for any other known TA system of this family. To further confirm the stoichiometry and assess the stability of the association, a 6:2 ParD2:ParE2 precursor of 17+ of charge was selected for MS/MS analysis. Acceleration of the ions was increased by applying higher voltages which lead to increased collisions that gradually caused the dissociation of the complex. New peaks corresponding to monomers of ParD2 were now detected in the spectra, with the concomitant detection of peaks corresponding to the mass of a 5:2 association between antitoxin and toxin (Fig. 1).

**Figure 1.**
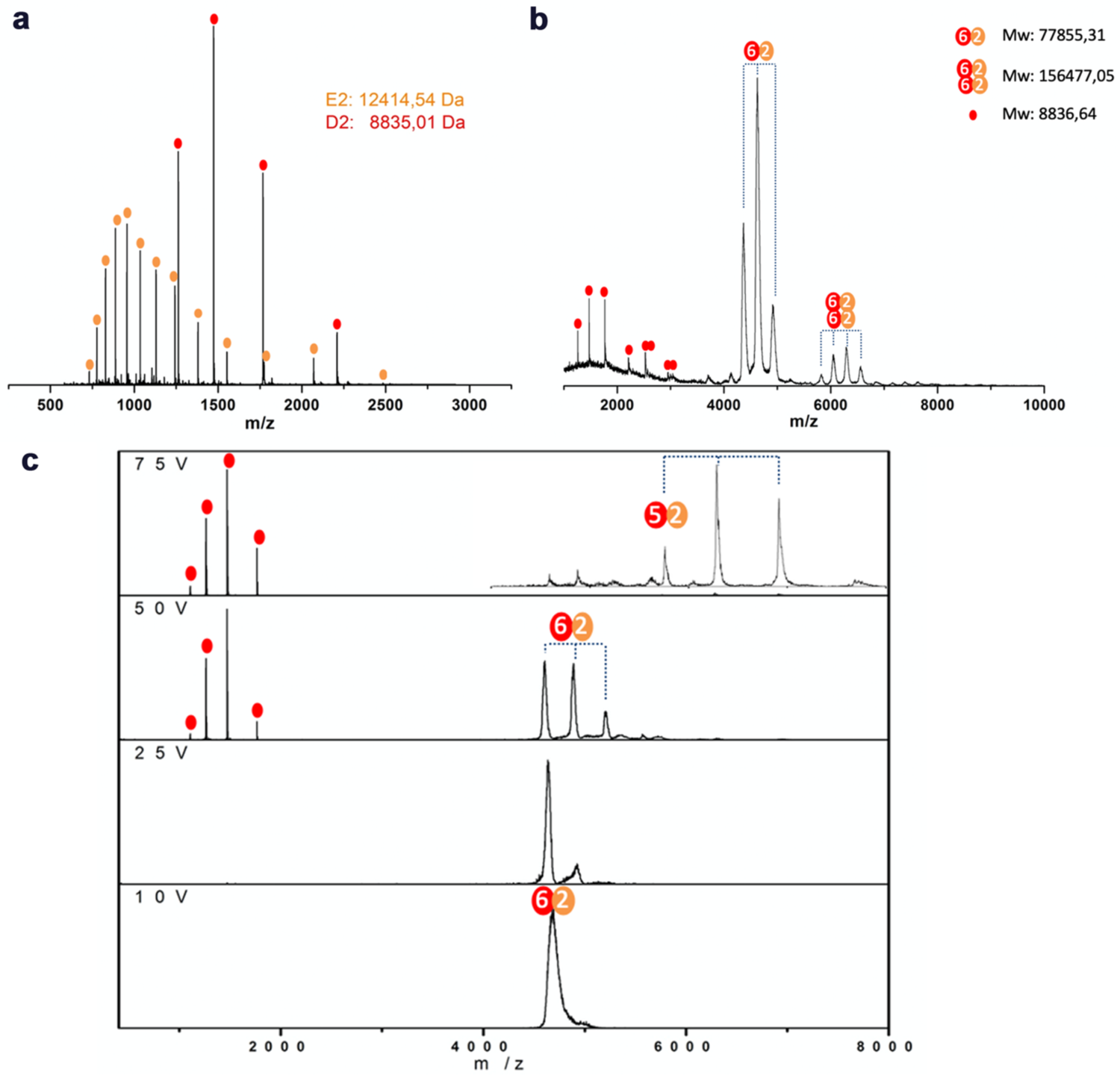
Mass spectrometry measurements with ParD2:ParE2 purified sample show a 6:2 TA stoichiometry. The measured and theoretical masses of the proteins and complexes are indicated. (a) Mass-spectrometry profile of the co-purified ParD2:ParE2 complex under denaturing conditions. All peaks were assigned to either ParE2 or ParD2 monomers, confirming the purity of the sample and the absence of degraded protein. (b) Native mass-spectrometry analysis of the co-purified ParD2:ParE2 complex. (c) MS/MS analysis of a 6:2 ParD2:ParE2 precursor (17+). Higher voltages in the trap-collision chamber were applied as indicated. Critical voltages and pressures used during the measurements were 25 V on the sampling cone, 10 V trap-collision energy for the sample in (b).

### Denaturant-induced dissociation of the ParD2:ParE2 complex yields isolated and re-folded toxin and antitoxin

In order to obtain the isolated proteins from the ParD2:ParE2 complex, a denaturant-induced dissociation protocol based on the Phd:Doc method described by (Sterckx et al., 2015), was optimized. This protocol allowed to obtain pure and refolded ParD2 and ParE2 proteins as confirmed by native MS and circular dichroism (CD) measurements (not shown), which were subsequently used for biophysical and DNA-binding experiments.

### ParD2:ParE2 complex exists as a 6:2 hetero-octamer in the crystal

In order to gain new insights into the mechanism of toxin inhibition by direct antitoxin interaction, the purified complex was crystalized in absence and presence of *Vc*ParD2:ParE2-specific camelid single-chain antibody fragments, which act as crystallization chaperones. Isolated *Vc*ParD2:ParE2 crystallized in space group P4_3_2_1_2 (96), and the structure was determined at 2.95 Å resolution. The asymmetric unit contains one ParE2 monomer and one full length ParD2 chain. ParD2 N-terminal domain is folded into the ribbon-helix-helix motif present in RK2 ParD. *Vc*ParD2 *α*-helix 2 extends over 48 Å, followed by a 7-residues long coil region connecting C-terminal *α*-helix 3. ParD2 interacts with the toxin monomer via the C-terminal half of *α*-helix 2, the following coil region and *α*-helix 3. The asymmetric unit additionally contains two other ParD2 molecules, for which no density is observed for their 30 C-terminal residues. The lack of electron density for the C-terminal tails of these two ParD2 chains stems from their intrinsically disordered nature and from not being contacted by toxin molecules due to the molar excess of ParD2 in the sample (Fig. 2a, Fig. 1). These two additional copies of ParD2 in the crystal confirm that the intrinsically disordered C-terminal ends of the antitoxin fold upon binding ParE2 monomers.

**Figure 2.**
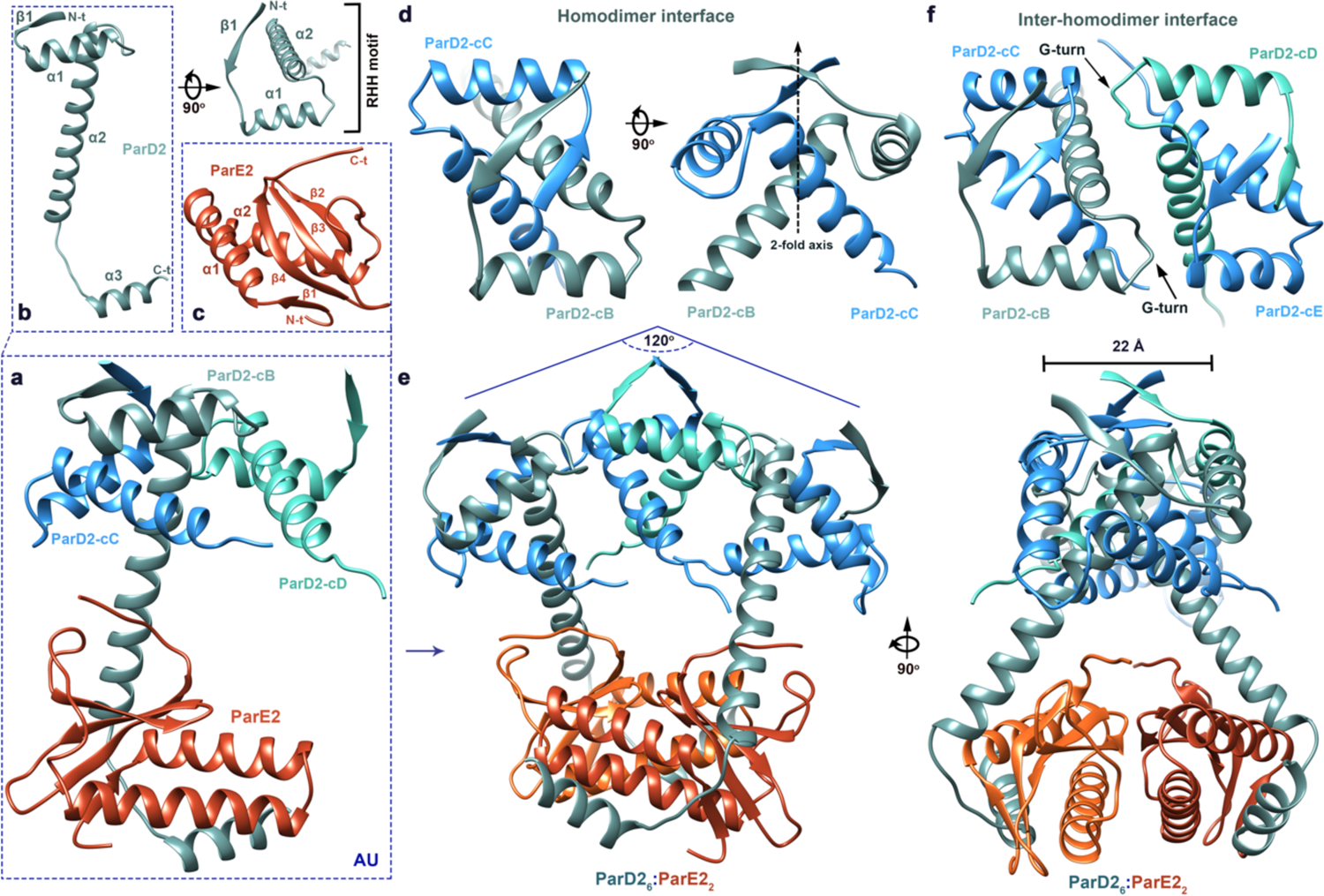
High resolution structure of the *Vc*ParD2:ParE2 complex. The disordered ParD2 C-terminus folds upon binding its toxin while the N-terminus adopts a RHH motif involved in homodimerization and DNA binding. (a) Asymmetric unit containing one full length ParD2 monomer interacting with ParE2 toxin via its C-terminal end. Ribbon representations of the fully folded (b) ParD2 antitoxin, with the secondary structure elements annotated along with the N and C termini and the RHH motif, and (c) ParE2 toxin. (d) Ribbon representation of the ParD2 homodimer. The 2-fold symmetric homodimerization interface involves the RHH motifs from both protomers and *α*-helices 2. (e) Ribbon representation of the 6:2 ParD2:ParE2 assembly generated by crystal symmetry. Three ParD2 homodimers interact via a second inter-homodimer interface to generate a 120° arch. The RHH motifs are located on the surface, which has a width of 22 Å, strongly suggesting this to be the DNA binding site. (f) Interactions involved in the ParD2 inter-homodimer interface include the highly conserved glycine-mediated turn between *α*-helices 1 and 2 in the CopG/Arc/MetJ family of repressors.

ParD2 C-terminus folds into an extended *α*-helix 2, followed by a coil region spanning G62-D67 connecting *α*2-*α*3. The interface between toxin and antitoxin buries a total area of 2707 Å^2^, which represents 22 % of ParE2 solvent-accessible area buried upon antitoxin binding. *Vc*ParD2 contacts with ParE2 monomer include electrostatic interactions between the negatively charged ParD2 C-terminus and positively charged regions on the surface of the toxin. Additionally, two hydrophobic patches on the surface of ParE2 are buried upon antitoxin binding (Fig. 3a). The first *Vc*ParE2 hydrophobic patch involves residues relatively conserved within the ParE family: F4 and L6 from *β*1, L14 and A18 in *α*1, L34 and F41 from *α*2. ParD2 C-terminal *α*-helix 3 buries several hydrophobic residues on this ParE2 patch, which include A63, Y65, L67, F70, I71 and L74 (Fig. 3c). The second hydrophobic stretch on ParE2 locates mostly on the *β*-sheet (F72, I84 and L87), with I57 and Y61 from the loop between *α*2 and *β*2 also involved in the interactions. ParD2 C-terminal end of *α*-helix 2 caps over this second patch involving a stretch of conserved leucines: L47, L50, L53 and L54 (Fig. 3b). Remarkably, the bottom of this second hydrophobic patch on ParE2 is lined with conserved F72, which contacts L50 and L53 from ParD2. This observation is reminiscent of the interactions between ParD and ParE in the *C. crescentus* complex (Dalton and Crosson, 2010).

**Figure 3.**
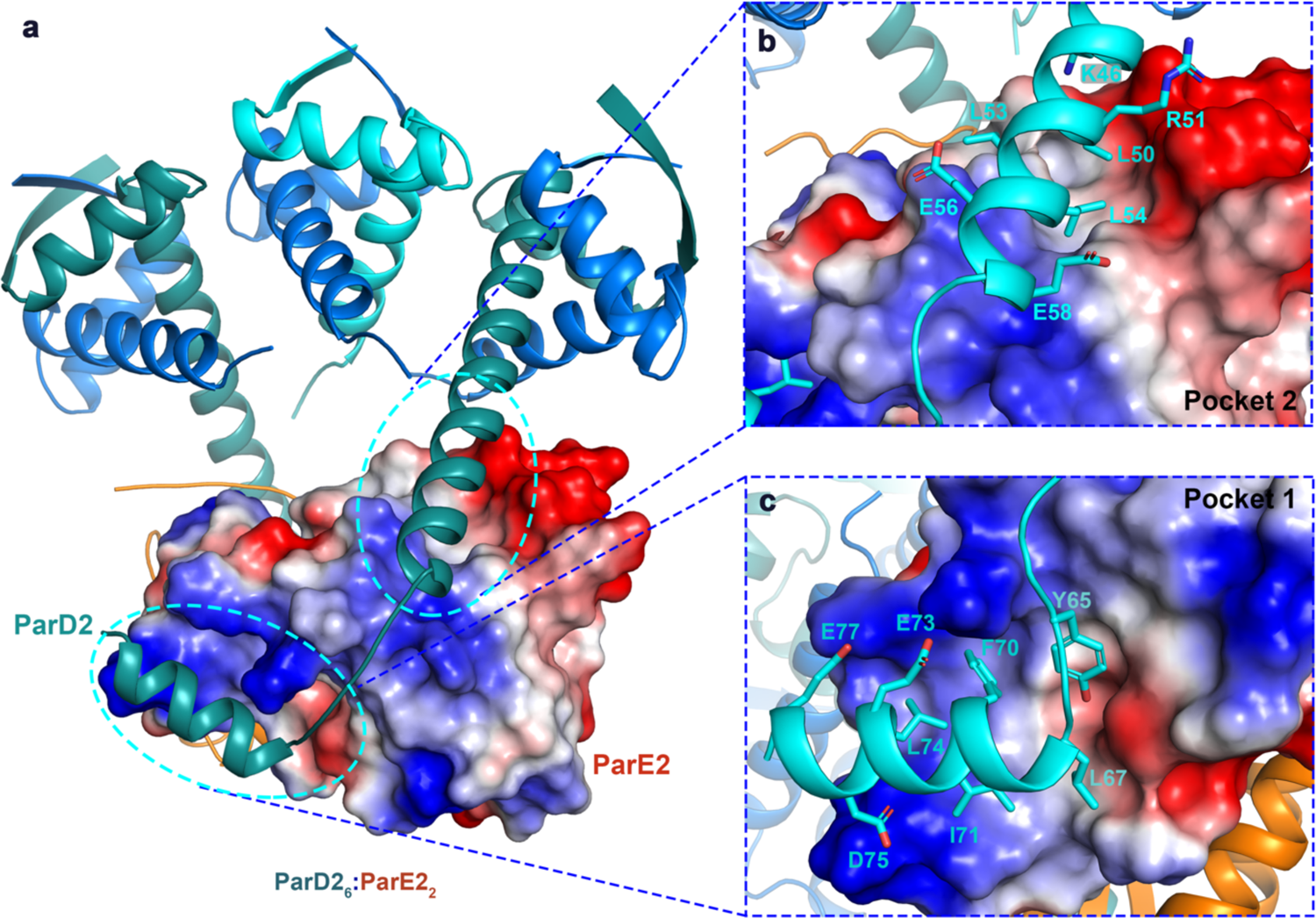
*Vc*ParD2 and *Vc*ParE2 interact via two hydrophobic patches surrounded by electrostatic interactions. (a) 6:2 ParD2:ParE2 crystallographic assembly. ParD2 chains are colored as in Fig. 2 and represented as ribbons. The surface of ParE2 is colored by electrostatic potential (red, negative and blue positive) as calculated in Pymol. The two main contact regions between toxin and antitoxin are encircled. (b) Hydrophobic patch 2 and (c) hydrophobic patch 1, as in the main text. ParD2 amino acid sidechains involved in interactions in this pocket are shown as sticks and labeled. ParD2 residues involved in salt-bridges are also represented as sticks.

The complex is additionally stabilized by several electrostatic interactions between the acidic ParD2 C-terminus and ParE2 basic regions. For example, surrounding hydrophobic region 1 on ParE2, residues K11, R15 and R30 are engaged in salt bridges with ParD2 residues on *α*3 E73, E77 and D75, respectively (Fig. 3c). Likewise, the coil region extending between ParD2 *α*2 and *α*3 is stabilized by electrostatic interactions, e.g., D62 on ParD2 is forming a salt bridge with R82 from ParE2. The C-end of ParD2 a2 is also engaged in some electrostatic interactions with ParE2 residues located on *β*4 and the C-terminal coil. For example, ParD2 E56 forms a salt bridge with R85 from ParE2 *β*4 (Fig. 3b). These interactions between ParD2 and ParE2 sequester the toxin thereby preventing its interaction with gyrase and effectively protecting the cell (Fig. 2b).

The N-terminal domains of ParD2 monomers adopt a ribbon-helix-helix DNA-binding motif. Two monomers associate into 2-fold symmetric dimers via the antiparallel juxtaposition of the N-terminal *β*-strands (Fig. 2d), which are presented to the surface of the dimer and constitute the DNA-binding site in this family of repressors (see below).

Structurally, ParE2 toxin belongs to the RelE family of toxins, although the activity diverged between these two proteins, the latter functioning as a ribosome-dependent mRNase. Despite the low sequence similarity shared by these proteins, the fold of ParE2 is equivalent to the one adopted by most members of this family: a four-stranded antiparallel *β*-sheet core flanked by two N-terminal *α*-helices in a helix-hairpin-helix motif mediated by a glycine short turn (Fig. 2a,c). The few ParE toxins structurally characterized so far, including *Vc*ParE2, have longer *α*-helices by approximately one extra turn, when compared to most RelE monomers (Fig. 4). A number of RelE-like RNases have been structurally characterized so far, however, ParE toxins are lacking in the databases. A DALI search identified *C. crescentus* ParE (RMSD of 1.4 Å over 92 C*α*, PDB ID 3kxe), *M. opportunistum* ParE (RMSD of 2.0 Å over 90 C*α* PDB ID 5ceg), *E. coli* O157:H7 ParE (RMSD of 1.9 Å over 85 C*α* PDB ID 5cw7), *E. coli* RelE (RMSD of 1.64 Å over 63 C*α* PDB ID 4fxe); *M. tuberculosis RelE3* (RMSD of 1.31 Å over 53 C*α* PDB ID 3oei) as the closest structural homologs to *Vc*ParE2.

**Figure 4.**
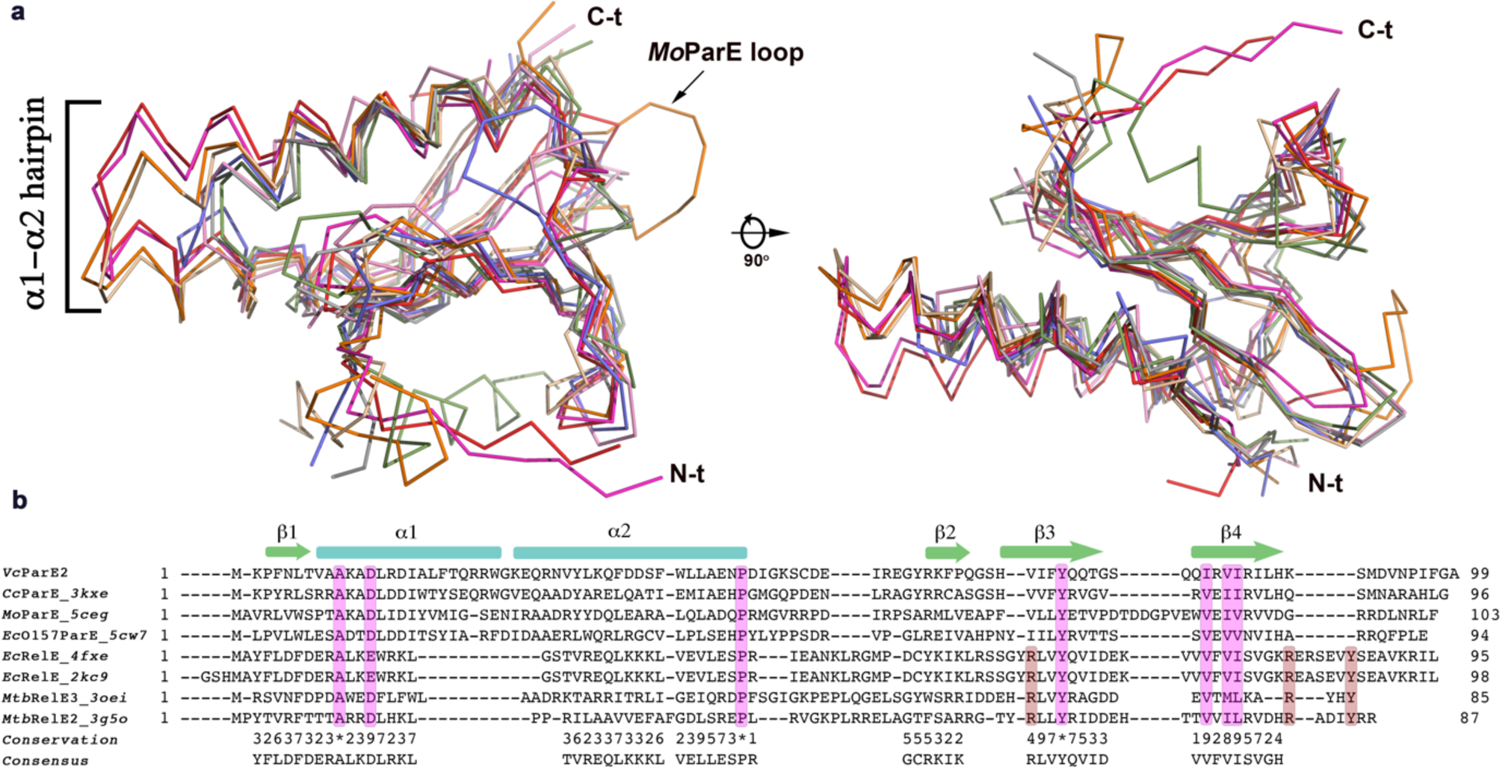
V. cholerae ParE2 adopts the RelE canonical fold, amidst some differences. (a) Ribbon representation of superimposed monomers of *Vc*ParE2, *V. cholerae* ParE2 (red); *Cc*ParE, *C. crescentus* ParE (magenta, PDB ID 3kxe); *Mo*ParE, *M. opportunistum* (orange, PDB ID 5ceg); *Ec*O157ParE, *E. coli* O157:H7 ParE (wheat, PDB ID 5cw7); *Ec*RelE, *E. coli* RelE (gray, PDB ID 4fxe); *Ec*RelE, *E. coli* RelE (light green, PDB ID 2kc9); *Mtb*RelE3, *M. tuberculosis* RelE3 (slate, PDB ID 3oei), *Mtb*RelE2, *M. tuberculosis* RelE2 (light pink, PDB ID 3g5o). The *α*1-*α*2 longer hairpin in the ParE family is labeled along with the long loop insertion in *M. opportunistum* ParE. (b) Structural alignment of the conserved fold in the RelE family. Highly conserved residues are highlighted in pink, while RelE catalytic residues are highlighted in red-brown.

Remarkably, the 6:2 ParD2:ParE2 stoichiometric organization previously observed in native mass spectrometry experiments could be generated by crystal symmetry. The assembly consists of a trimer of ParD2 N-terminal dimers, which associate via a second ParD2 inter-homodimer interface, forming an arch of approximately 120° (Fig. 2e,f). The angled surface formed by the three consecutive ParD2 dimers displays a width of approximately 22 Å, which taken together with the presence of the antiparallel *β*-sheets positioned on this surface, strongly suggests this to constitute the DNA-binding site (Fig. 2e). Two ParD2 molecules from the flanking homodimers extend *α*-helices 2 approximately 30° outward on either side of the central plane of the ParD2 homohexameric arch, trapping two ParE2 monomers in between (Fig. 2e). The interface between the two ParE2 monomers extends only over 850 Å^2^ on each monomer. Contacts are established between the *β*2-*β*3 loops from both monomers, and the end of the *α*-hairpins and the long *α*2-*β*2 loop (Fig. 2e).

### VcParD2:ParE2 complex is recognized as a 6:2 complex by complex-specific nanobodies

Nanobodies were generated against the ParD2:ParE2 complex, and crystals were screened with three of the best binders. Crystals grew in presence of Nb80 and data were collected up to 2.91 Å resolution. ParD2:ParE2:Nb80 crystallized in space group C2 (5). Importantly, the asymmetric unit contains the same 6:2 ParD2:ParE2 assembly seen in the ParD2:ParE2 crystal structure in absence of nanobody (RMSD of 0.305 Å over 528 Ca) (Fig 5a). Three nanobodies are bound to an epitope located on the 6:2 ParD2:ParE2 assembly itself, involving the two ParE2 protomers in close proximity and the C-terminal D66-S76 a-helix in ParD2 in complex with the toxin (Fig. 5b).

**Figure 5.**
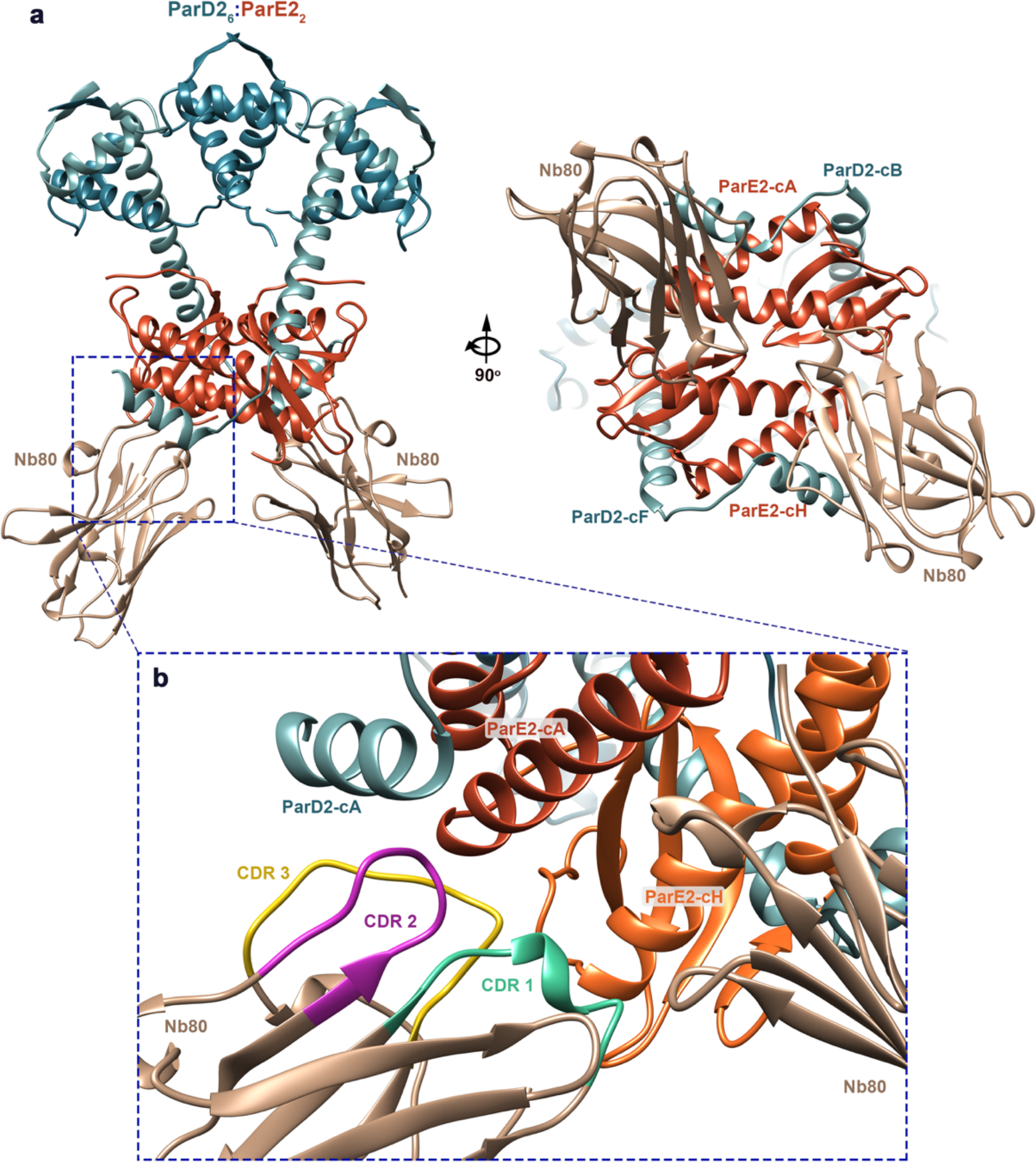
Nanobody 80 specifically recognized the 6:2 ParD2:ParE2 assembly. (a) Ribbon representation of the 6:2 ParD2:ParE2 assembly in the asymmetric unit of ParD2:ParE2-Nb80 crystals. Nanobody molecules are bound to an epitope composed by two ParE2 protomers and ParD2 C-terminal *α*-helix 3. (b) Ribbon representation of Nb80 bound to the epitope on ParD2:ParE2. The CDR regions are highlighted in different colors.

### The crystallographic 6:2 ParD2:ParE2 hetero-octameric assembly is present in solution

In order to further assess the stoichiometry of the ParD2:ParE2 complex in solution, SAXS experiments with pure complex were performed. The Porod volume estimation of the molecular weight of the ParD2:ParE2 complex in solution shows a value of 77.11 kDa, which is practically the same as the expected theoretical value for the 6:2 TA association (78.22 kDa). These results further agree with the analytical gel filtration and native mass spectrometry experiments above. In order to determine whether the crystallographic assembly exists in solution, the missing C-termini of the four ParD2 protomers were modelled, including all missing sidechains and loops, to be able to compare the theoretical scattering profile with the experimental data. This process yielded an all-atom ensemble of models that agrees with the crystallographic data with *χ^2^* value of 1.09, therefore confirming the presence of the crystallographic assembly in solution (Fig. 6).

**Figure 6.**
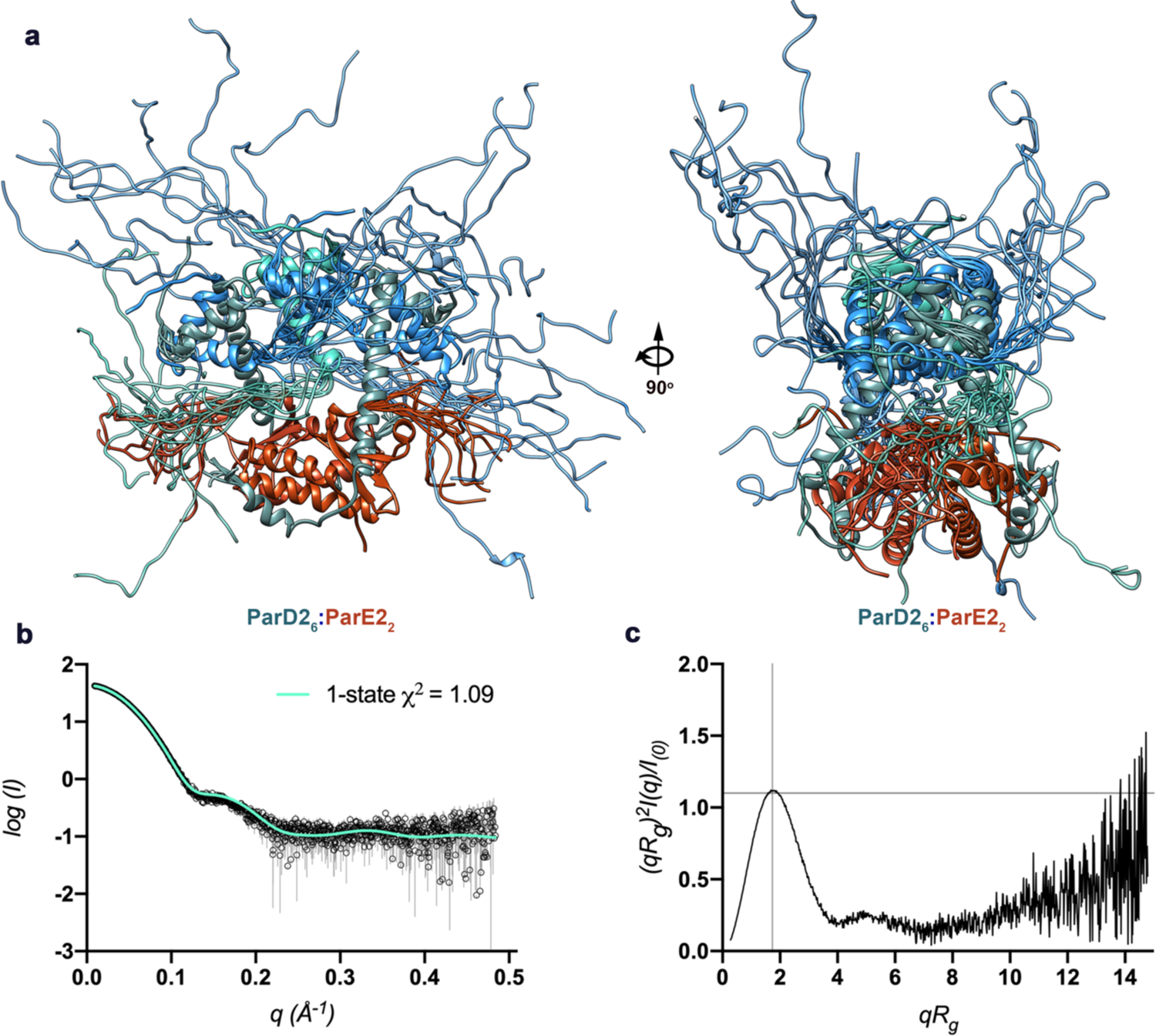
Free ParD2 C-termini are intrinsically disordered. (a) Ribbon representation of the 10 best SAXS models of the 6:2 ParD2:ParE2 assembly. (b) Solution scattering curve of the ParD2:ParE2 complex. The theoretical profile for the ensemble shown in (a) is compared to the experimental data (*χ^2^* = 1.09). (c) Kratky plot for ParD2:ParE2 6:2 complex showing the characteristic features of a globular particle in solution.

Additionally, the dimensionless Kratky plot for the ParD2:ParE2 complex shows a maximum around the expected values for a globular particle (1.73; 1.1), which is not surprising given that the generated ensemble of all-atom models shows the missing C-termini to be stacking up on both sides of the folded core, which confers an overall globular shape to the particle (Fig. 6c).

### ParD2 antitoxin forms 2-fold symmetric dimers via the N-terminal RHH motifs

ParD2 homodimerizes via the conserved ribbon-helix-helix (RHH) DNA binding motif also found in other ParD antitoxins (plasmid RK2 ParD, PDB ID 2an7 and *C. crescentus* ParD, PDB ID 3kxe). This topology consists of four helices in an open array of two hairpins often found in bacterial and phage repressors. Additionally, inspection of the electrostatic surface of the dimer of ParD2 reveals a positively charged area comprising the N-terminal b-ribbons, which constitutes the DNA binding region in this type of proteins. Interestingly, ParD2 homodimers associate into higher order oligomers via the opposite side of the hairpins interface, through the aforementioned inter-homodimer interface (see Fig. 2f).

### ParD2 belongs to the CopG/MetJ/Arc family of transcriptional repressors

The N-terminus of the *Vc*ParD2 monomer superposes with RMSD of 2.21 Å over 31 C*α* pairs of RK2 ParD (Oberer et al., 2007, PDB ID 2an7) and with RMSD of 0.54 Å over 42 C*α* atoms of *C. crescentus* ParD (Dalton et al., 2010, PDB ID 3kxe). In a DALI search, many structural homologs from this family showed similarity to *Vc*ParD2, like the recently characterized CopA antitoxin (Zhao et al., 2019; PDB ID 6iya), which is part of a newly identified type II TA module in *Shewanella oneidensis*. The RMSD for the alignment over 43 C*α* atoms of CopA and ParD2 has a value of 1.8 Å with a DALI z-score of 5.6. It also shows structural homology to the N-terminal domain of antitoxin AtaR (DUF1778-domain containing, 2.1 Å RMSD over 44 C*α*, 5.2 z-score) of the AtaT-AtaR TA pair (Yashiro et al., 2019; PDB ID 6ajn).

Remarkably, the 45-residue long transcriptional repressor CopG (Gomis-Rüth et al., 1998; PDB ID 2cpg), which is one of the archetypes of this family, shows a 1.8 Å RMSD value over 41 C*α* pairs of the N-terminus of *Vc*ParD2 monomer. Moreover, the dimerization of CopG highly resembles that of *Vc*ParD2, showing an RMSD value of 2.8 Å over 76 C*α* of the dimer. The DALI search output parameters are summarized in Table 2. The dimeric form adopted by *Vc*ParD2 structural homologs is highly similar, with the symmetrical placement of both chains via a local two-fold axis and the concomitant formation of the N-terminal antiparallel *β*-strand (Fig. 7).

**Figure 7.**
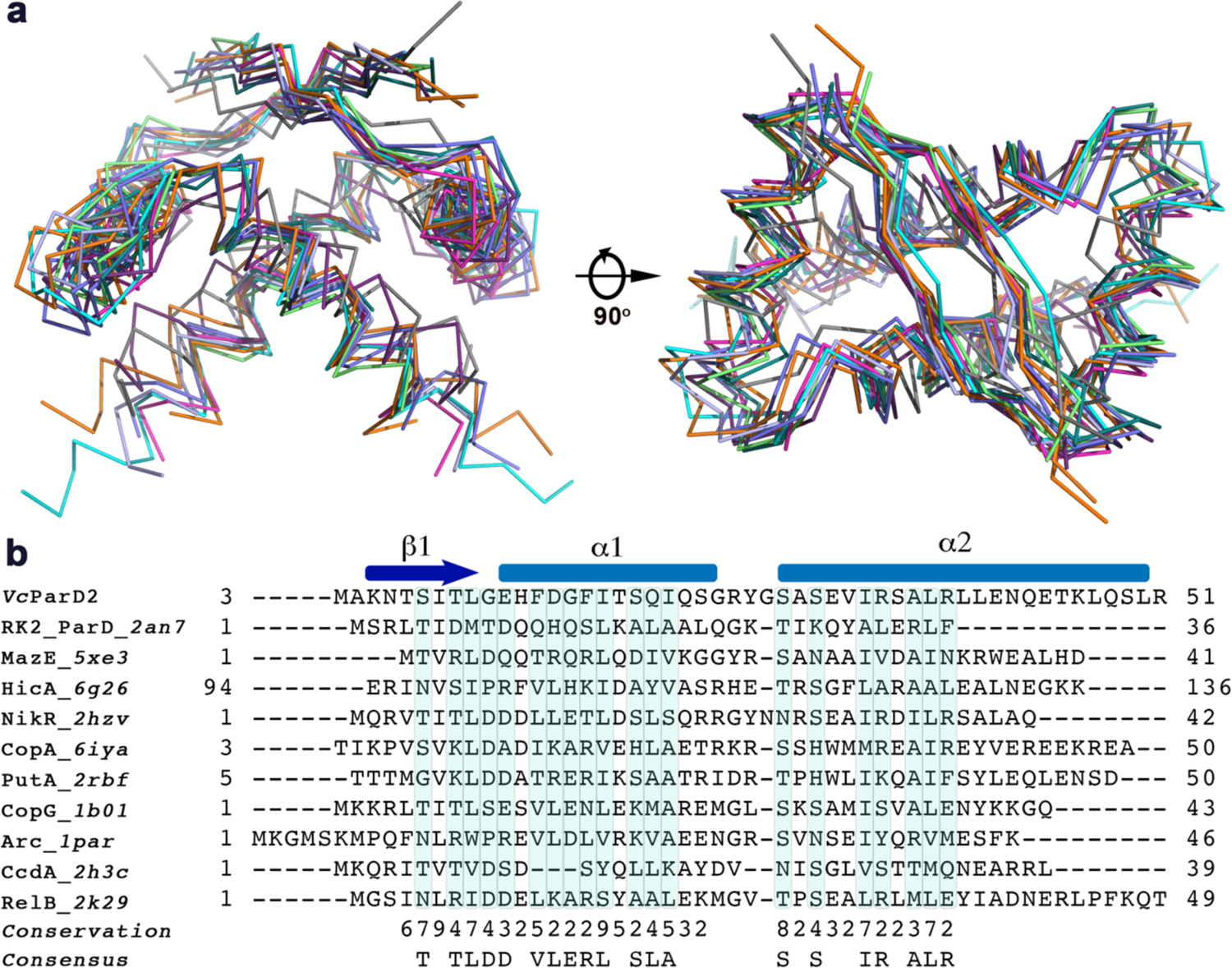
V. cholerae ParD2 belongs to the CopG/Arc family of transcriptional repressors. (a) Ribbon representation of the superposition of RHH dimers from *Vc*ParD2, *V. cholerae* ParD2 antitoxin (cyan); RK2_ParD, *E. coli* RK2 ParD antitoxin (dark green, PDB ID 2an7), MazE_5xe3, *E. coli* MazE antitoxin (magenta, PDB ID 5xe3); HicA, *B. pseudomallei* HicA antitoxin (light purple, PDB ID 6g26); NikR, *E. coli* NikR repressor (slate blue, PDB ID 2hzv); CopA, *S. oneidensis* CopA antitoxin (orange, PDB ID 6iya); PutA, *E. coli* proline utilization A (PutA) flavoprotein (blue, PDB ID 2rbf); CopG, *S. agalactiae* CopG repressor (dark orange, PDB ID 1b01); Arc, *Salmonella* virus P22 Arc repressor (green, PDB ID 1par); CcdA, *E. coli* CcdA antitoxin (gray, PDB ID 2h3c); RelB, *E. coli* RelB antitoxin (dark purple, PDB ID 2k29).

**Table 1.** Crystallographic data collection and refinement statistics.

|  | **ParD2:ParE2 Complex\*** | **ParD2:ParE2-Nb80** |
| --- | --- | --- |
| **Wavelength** | 0.97 | 0.98 |
| **Resolution range** | 45.77 - 2.95 (3.06 - 2.95) | 46.55 - 2.91 (3.014 - 2.91) |
| **Space group** | P 4<sub>3</sub> 2<sub>1</sub> 2 | C 1 2 1 |
| **Unit cell** | 65.78 65.78 127.44 90 90 90 | 146.91 113.64 127.31 90 119.23 90 |
| **Total reflections** | 78228 (7628) | 273685 (24730) |
| **Unique reflections** | 6339 (605) | 40069 (3901) |
| **Multiplicity** | 12.3 (12.6) | 6.8 (6.3) |
| **Completeness (%)** | 99.81 (99.02) | 99.59 (98.98) |
| **Mean I/sigma(I)** | 10.62 (1.19) | 9.26 (1.22) |
| **Wilson B-factor** | 87.44 | 86.07 |
| **R-merge** | 0.16 (1.99) | 0.181 (1.39) |
| **R-meas** | 0.17 (2.08) | 0.19 (1.51) |
| **R-pim** | 0.048 (0.58) | 0.07 (0.58) |
| **CC1/2** | 0.99 (0.512) | 0.99 (0.67) |
| **CC\*** | 1 (0.82) | 0.99 (0.89) |
| **Reflections used in refinement** | 6339 (605) | 39986 (3896) |
| **Reflections used for R-free** | 317 (30) | 1997 (195) |
| **R-work** | 0.24 (0.37) | 0.23 (0.45) |
| **R-free** | 0.29 (0.47) | 0.28 (0.47) |
| **CC(work)** | 0.95 (0.63) | 0.94 (0.73) |
| **CC(free)** | 0.91 (0.28) | 0.94 (0.66) |
| **Number of non-hydrogen atoms** | 2046 | 8474 |
| **macromolecules** | 2046 | 8474 |
| **Protein residues** | 264 | 1130 |
| **RMS(bonds)** | 0.006 | 0.005 |
| **RMS(angles)** | 0.91 | 0.96 |
| **Ramachandran favored (%)** | 91.80 | 91.32 |
| **Ramachandran allowed (%)** | 8.20 | 7.50 |
| **Ramachandran outliers (%)** | 0.00 | 1.19 |
| **Rotamer outliers (%)** | 0.00 | 0.00 |
| **Clashscore** | 14.08 | 13.42 |
| **Average B-factor** | 106.16 | 102.19 |
| **macromolecules** | 106.16 | 102.19 |
| **Number of TLS groups** | 18 | 72 |

Statistics for the highest-resolution shell are shown in parentheses.

**Table 2.**
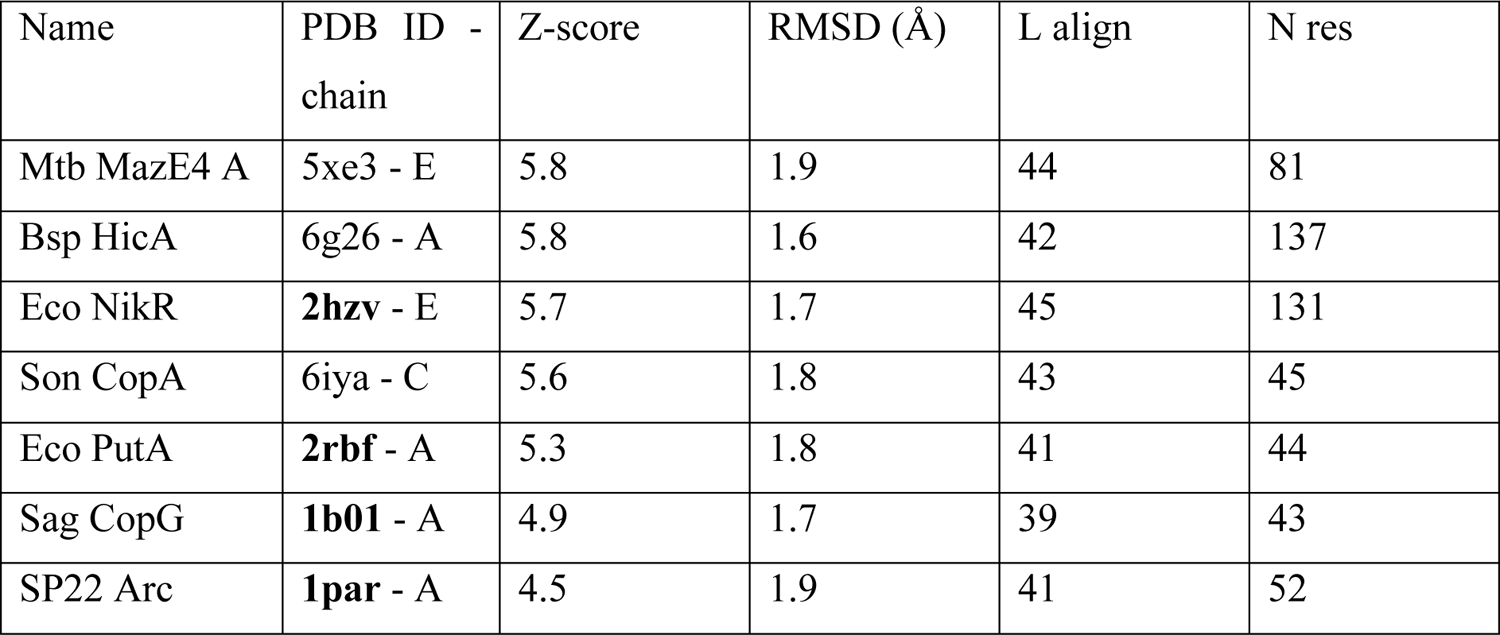
DALI results for *Vc*ParD2

### Isolated ParD2 and the ParD2:ParE2 complex bind the parDE2 promoter/operator region

EMSAs were used to study binding of the purified ParD2:ParE2 complex, ParD2 antitoxin and ParE2 toxin to a [5’-^32^P] single-end labelled 151 bp-long DNA fragment corresponding to the sequence upstream of *parD2* (−60 to +91 with respect to the BPROM predicted start of transcription) (Fig. 8). A wide range of protein concentrations was used in all cases, keeping the DNA concentration constant, to compare across proteins. The results indicate that both isolated ParD2 antitoxin and copurified ParD2:ParE2 complex are able to bind the DNA fragment. Three concentration-dependent ParD2:ParE2-DNA complexes with different migration velocities are observed (DE1 to DE3; Fig. 8a). When ParD2:ParE2 concentrations surpass 4 μmol l^-1^ (expressed in 6:2 ParD2:ParE2 stoichiometry equivalents), only one band is detected that hardly penetrates the gel due to precipitation. This observation suggests the concentration dependent formation of ParD2:ParE2-DNA complexes with a different stoichiometry and/or DNA conformation (e.g., bending, kinking, wrapping). On the other hand, isolated ParD2 antitoxin binds the 151 nt-long DNA fragment with less variability (Fig. 8b), resulting in two distinct and specific concentration dependent ParD2-DNA complexes with fast migration velocity (D1 and D2, Fig. 8b). When more than 6 μmol l^-1^ of ParD2 (in dimer equivalents) is incubated with DNA, only one retarded band with migration velocity similar or identical to that of D2 is visible. Strikingly, D1 and D2 show practically the same shift as DE1 and DE2, respectively (D2 and DE2 migrate together with ssDNA while D1 and DE1 are just below the latter). This could be explained by the antitoxin excess in the complex, which dissociates and exists as homodimer, as seen in native MS measurements. These free ParD2 homodimers could readily bind DNA as one homodimer-DNA complex (D1 and DE1) or two homodimers bound to two distal operator sites (D2 and DE2). As expected, isolated ParE2 toxin did not bind DNA (Fig. 8b). However, when isolated ParE2 was added to ParD2 antitoxin prior to incubation with DNA, the binding of ParD2 to the DNA was heavily altered. The supershift observed with the reconstituted ParD2:ParE2 complex incubated with DNA resembled the one previously observed with copurified ParD2:ParE2 (DE3). It is noteworthy that some precipitation in the wells is visible with the reconstituted complex and isolated ParE2 (Fig. 8b), which could have potentially arisen from the solution conditions of isolated ParE2. The specificity of binding of copurified ParD2:ParE2 complex and isolated ParD2 antitoxin to the labelled 151 nt-long DNA fragment was further investigated by the addition of non-specific competitor DNA (non-labelled sonicated salmon sperm DNA) (Fig. 8c,d). The addition of 0.0125 mg ml^-1^ non-specific competitor in ParD2:ParE2 binding assays caused slight smearing of the bands, particularly those with lower mobility, which could indicate the presence of some level of unspecific binding in these most retarded complexes. However, binding proved to be specific, since even at a very large excess of competitor DNA (0.16 mg ml^-1^, approximately 2000-fold excess), the formation of the fastest migrating complexes was still observed, and all previously identified species (DE1-DE3) were still visible. The formation of ParD2:ParE2-DNA complexes only started to decrease at very high concentrations of non-specific competitor (0.64 mg ml^-1^), and completely vanished at 2.56 mg ml^-1^. On the other hand, ParD2-DNA binding was seemingly not affected by 0.0125 mg ml^-1^ of non-specific competitor DNA (Fig. 8d), moreover, partial inhibition of ParD2-operator DNA complex formation was only observed at very high concentrations (2.56 mg ml^-1^) of non-specific competitor DNA (Fig. 8d). These results indicate that the binding of ParD2 to the operator is highly specific.

**Figure 8.**
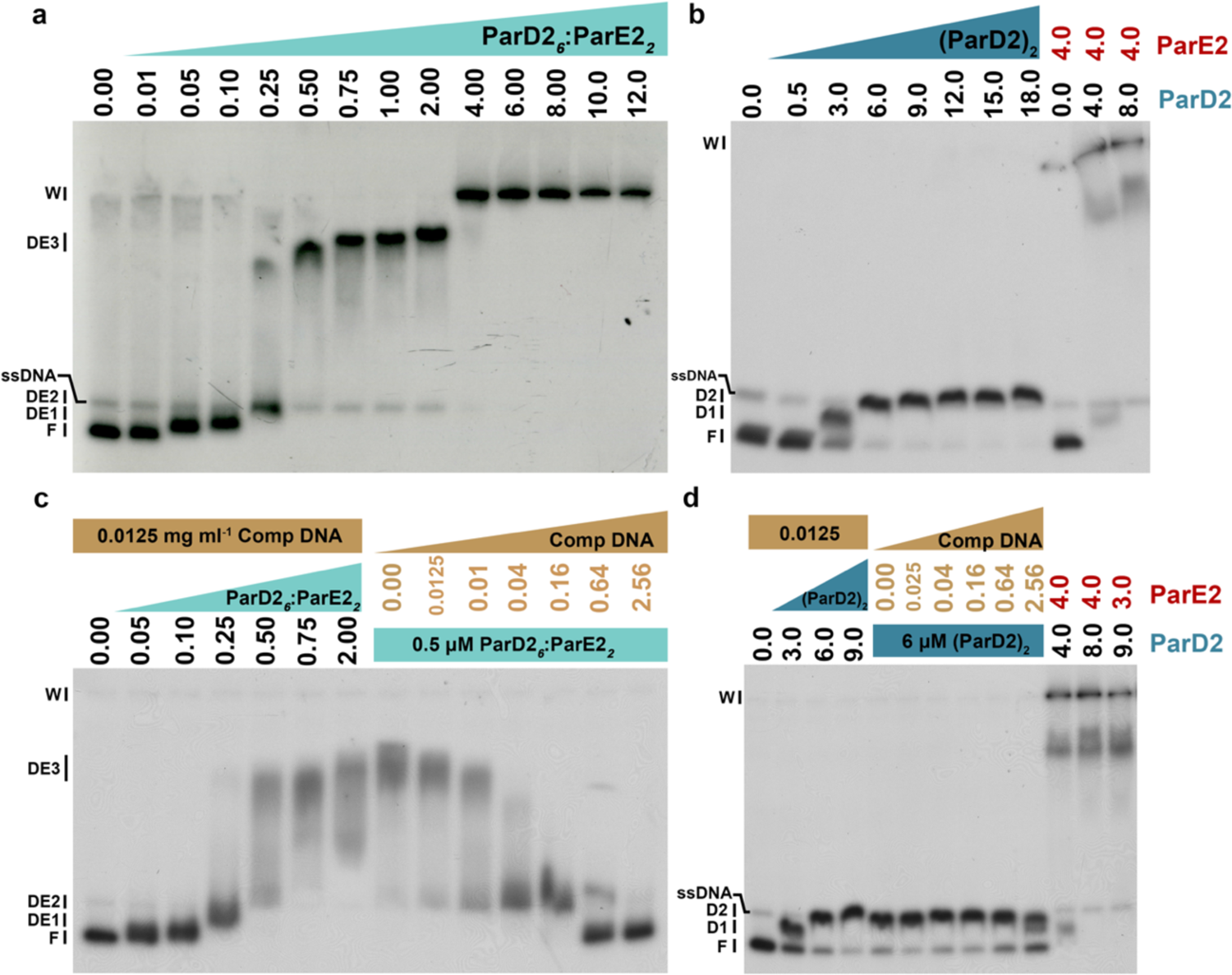
Representative autoradiographs of EMSAs with binding of various proteins to a [5’-*^32^*P] single-end labelled 151 bp DNA fragment, extending from position −60 to +91 with respect to the transcription start of the *parDE2* operon. (a) Binding of increasing concentrations (indicated in µM) of copurified 6:2 ParD2:ParE2 complex. The position of free DNA (F), single stranded DNA (ssDNA), the bottom of wells (W), and of protein-DNA complexes (DE1 to DE3) with different migration velocities are indicated. (b) Binding of increasing concentrations of isolated ParD2 (in µM of dimer equivalent) and of isolated ParD2 preincubated with ParE2 in different stoichiometries. (c) Binding of increasing concentrations of isolated 6:2 ParD2:ParE2 (in µM) in presence of a constant amount of non-specific non-labelled competitor DNA (sonicated salmon sperm DNA) and constant concentration of 6:2 ParD2:ParE2 assembly in the presence of increasing concentrations of non-specific competitor DNA (in mg ml^-1^), as indicated. (d) Binding of increasing concentrations of ParD2 (in µM of dimer equivalents) in presence of a constant amount of non-specific competitor DNA and of a constant concentration of ParD2 (in dimer equivalents) in the presence of increasing concentrations of non-specific competitor DNA (in mg ml^-1^), as indicated, and of ParD2 preincubated with isolated ParE2 at different ratios.

### ParE2 toxin influences ParD2 DNA binding activity in a T:A ratio-dependent manner

To further assess the effects of changes in the stoichiometry of ParD2:ParE2 complex on DNA binding, titrations with isolated ParE2 were performed. In these assays, 0.5 and 0.25 μmol l^-1^ of copurified ParD2:ParE2 complex (equivalent to 3 and 1.5 μmol l^-1^ of ParD2, in 6:2 equivalents of complex stoichiometry), were titrated with isolated ParE2 in the presence of 0.0125 mg ml^-1^ of non-specific competitor DNA to reduce unspecific binding (Fig. 9). These data showed no DNA precipitation in the well and the formation of the same low mobility ParD2:ParE2-DNA complexes seen in the previous assay (Fig. 9a and Fig. 9c). Strikingly, when the ParE2:ParD2 ratio surpassed 1:1, DNA started to be released from protein-DNA complexes, and much faster migrating bands appeared, very close to the position of free DNA (Fig. 9a). This finding is reminiscent of the conditional cooperativity effect observed for other TA systems (Overgaard et al., 2008; Garcia-Pino et al., 2010) and the effect of RK2 ParE on ParD DNA binding (Johnson et al., 1996). The assay was repeated at higher ParE2:ParD2 ratios (Fig. 2b) but lower ParD2 concentrations in order to prevent precipitation as observed before. At lower total protein concentration most of the DNA is visible as the DE2 species (along with some smearing), as seen in the EMSAs with the ParD2:ParE2 complex (Fig. 8a). Strikingly, initial increases in toxin:antitoxin ratios abrogated DE3 and other slower migrating species, with the appearance of faster migrating bands corresponding to DE2 and DE1 (Fig. 9). When the ParE2:ParD2 ratio exceeded 1, a band corresponding to the position of free DNA became visible (Fig. 9b). Overall, the data demonstrate that ParE2 affects the DNA binding of ParD2 by inducing the formation of low mobility complexes (supershifted) when the ParE2:ParD2 ratio remains below 1. However, when ParE2 concentrations increase over those of available ParD2, ParD2-DNA binding is affected, and DNA release occurs.

**Figure 9.**
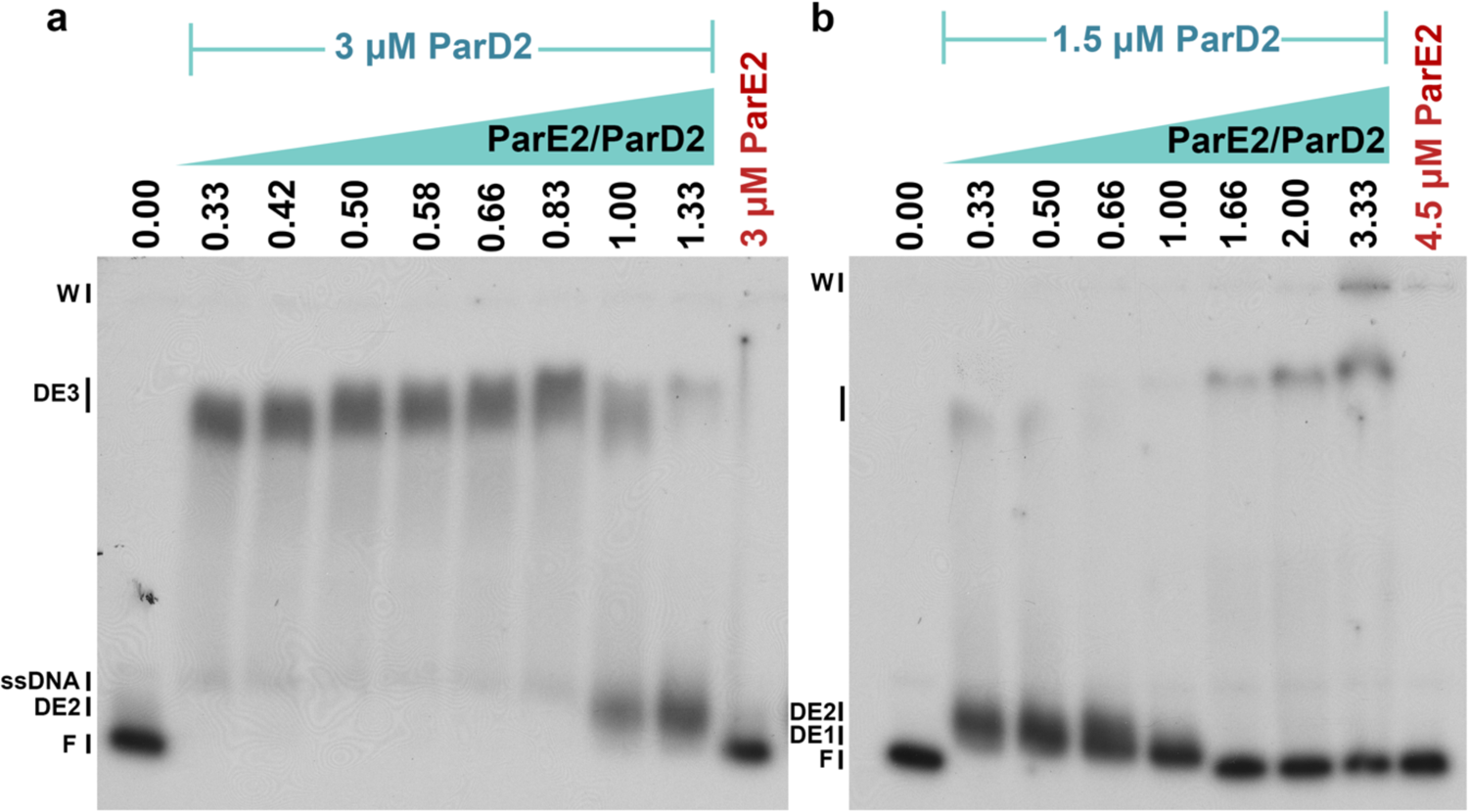
ParE2 influences ParD2-DNA binding in a toxin/antitoxin ratio dependent manner. Representative autoradiographs of EMSAs with binding of a constant amount of copurified 6:2 ParD2:ParE2 corresponding to 3 µM ParD2 (a) or 1.5 µM (b), preincubated with increasing amounts of isolated ParE2 resulting in different ParE2/ParD2 ratios as indicated, prior to incubation with the single-end labelled 151 bp DNA fragment comprising the control region of the *parDE2* operon.

### Sequence-specific ParD2-DNA interactions

In order to delimit the ParD2 binding site in the *parDE2* control region, DNase I footprinting assays were performed with isolated ParD2 antitoxin and copurified ParD2:ParE2 complex binding to the single-end labelled 151 bp-long control region. Surprisingly, ParD2 DNase I footprinting did not show any protected region on the DNA, not even at high ParD2 concentrations (96 μmol l^-1^; results not shown). However, when the copurified ParD2:ParE2 complex was used, DNase I footprinting revealed an approximately 36 nt-long zone of protection, from position −21 to +13 on the top (coding) strand and from −22 to +10 on the bottom (template) strand (Fig. 10a,b). A clear protection of ParD2:ParE2 binding against DNase I was visible at all protein concentrations, except at the lowest value of 0.1 μmol l^-1^. Furthermore, hyperreactivity for DNase I was detected on both strands, near the extremities of the protected regions, which suggests local ParD2:ParE2 induced DNA bending (minor groove widening). Remarkably, this hyperreactivity is most pronounced at the lowest ParD2:ParE2 concentrations, even when protection of the other bands is still rather weak. Hyperreactivity decreases as ParD2:ParE2 concentrations and protection increases (bands become fainter), without further extension of protected area size. A possible explanation for the decrease in hyperreactivity could be the introduction of relaxation of local DNA bending at the specific binding site-proximal ends as protein concentration increases caused by unspecific binding on distal sites.

**Figure 10.**
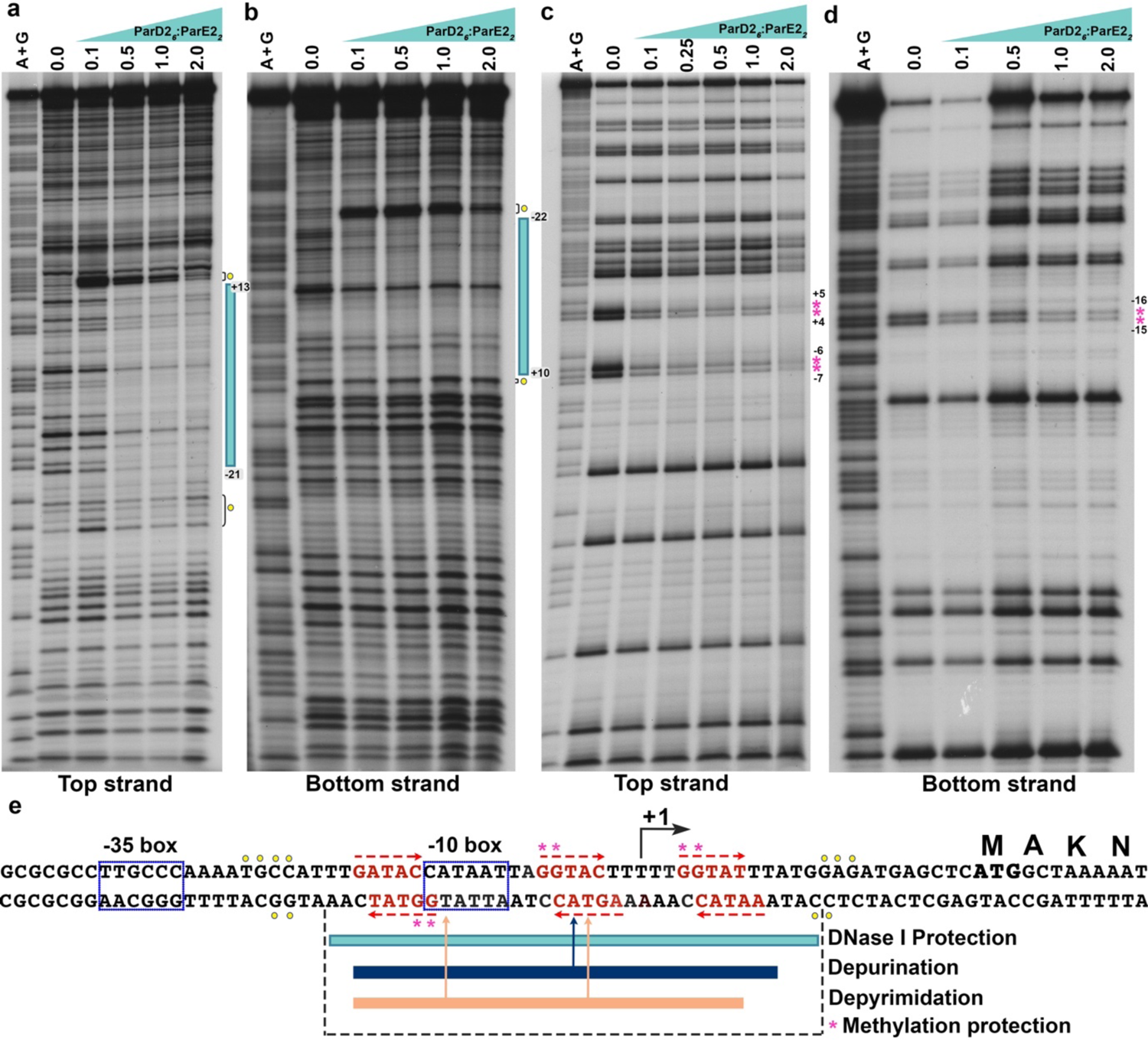
DNase I footprinting and methylation protection. (a,b) Autoradiographs of DNase I footprinting with increasing concentrations of copurified 6:2 ParD2:ParE2 (in µM) to the 151 bp fragment with either the top (coding) or bottom (template) strand labelled. Filled vertical bars indicate the regions of protection. Positions of hyperreactivity are indicated with a filled yellow circle. A+G corresponds to the Maxam-Gilbert sequencing ladder. (c,d) Methylation protection. Aliquots of the [5’-^32^P] single-end labelled 151 bp parDE2 control region were incubated with various concentrations of ParD26:ParE22 (in µM), then sparingly modified with DMS, precipitated, cleaved at methylated guanine residues with piperidine and the reaction products analyzed by gel electrophoresis in denaturing conditions. Guanine residues that are protected against methylation upon protein binding are indicated with an asterisk. A+G corresponds to the Maxam-Gilbert sequencing ladder. (e) Sequence of the *parDE2* control region. +1 indicates the start of transcription, the putative −10 and −35 promoter elements are boxed. Arrows indicate imperfect inverted repeats. The regions protected against cleavage by DNAse I and regions that upon base specific modification negatively interfere with complex formation (see Fig. 13) are indicated with a bar.

Additional information on critical base and groove-specific contacts was gathered by pre-modification binding interference (missing contact probing; Brunelle and Schleif, 1987) and purine methylation protection experiments. To reveal base specific contacts that contribute to the energy of complex formation, the 151 bp operator DNA was sparingly modified (on average one modification per molecule) with citrate (depurination) or hydrazine (depyrimidation) prior to incubation with ParD2:ParE2 at various concentrations, and complexes separated from free DNA by EMSA. Bands corresponding to different complexes were recovered together with free DNA and input control DNA (no protein added) from the EMSA gels, cleaved at modified positions by piperidine treatment at 90°C, and the reaction products analyzed by denaturing gel electrophoresis (Fig. 11). Removal of a base will reduce the affinity of the modified DNA molecule for the protein if that base is important for DNA binding. As a consequence, this type of molecules will be underrepresented in the bound forms and overrepresented in the free form or smeared out between the positions of the bound and free forms, if enhanced dissociation occurs during migration. It is worth to note that the effects of base removal on protein binding can be direct (lack of a base-specific contact) or indirect (e.g., effects on local DNA flexibility). Premodification experiments were performed with both top strand and bottom strand labelled DNA fragments to detect any strand-specific effects. Complexes DE1 to DE4 were recovered from EMSAs and analyzed for interference effects. On the top strand, removal of any purine residue in the stretch from −20 to +7 resulted in a negative effect on the formation of complex DE3 (Fig. 11a,b), whereas, in contrast, no negative effects were observed on the formation of complex DE2. On the bottom strand, removal of any purine residue in the stretch from position −18 to +9 had a negative effect on the formation of complexes DE1 and DE2 (Fig. 11a,b). However, the most striking negative effect is observed upon removal of the adenine residue at position −5. The negative interference observed for the bound form of complex DE1 was accompanied by the overrepresentation of the corresponding bands in the free form (Fig. 11b). Notice that free DNA could only be recovered from these lanes as the necessity for higher protein concentrations to form the slower migrating complexes results in saturation of DNA binding. Interference effects on the formation of other complexes with still lower migration velocity was not analyzed further due to the low signal intensity and intense smearing of the bands.

**Figure 11.**
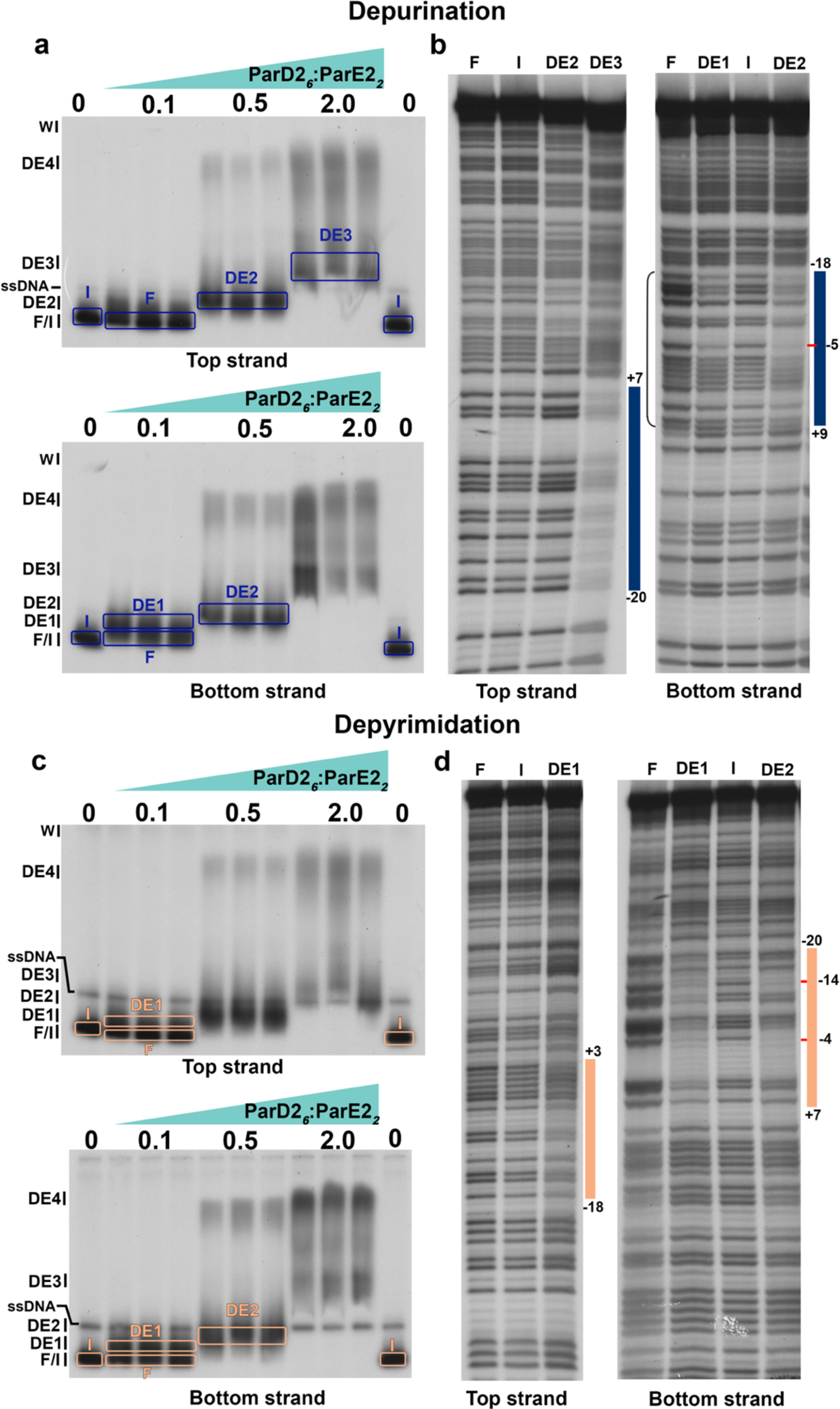
Identification of base specific contacts. (a to d) Missing contact probing with binding of copurified 6:2 ParD2:ParE2 at various concentrations to sparingly depurinated (a, b) or depyrimidated (c, d) 151 bp *parDE2* control region. Input DNA (I, no protein added), free DNA (F) and complexes with a different migration velocity were recovered from the EMSA gels, cleaved at modified positions with piperidine, and the reaction products separated by gel electrophoresis in denaturing conditions. Positions that upon modification result in reduced affinity for the protein complex are indicated with a blue (depurination) or orange (depyrimidation) vertical bar. The strongest negative signals are indicated with a short horizontal line.

Similarly, depyrimidation binding interference assays revealed weak interference effects for complex DE1 in the top strand, covering positions −18 to +3 (Fig. 11c,d). On the bottom strand, depyrimidation of any residue in the stretch extending from position −20 to +7 resulted in a negative effect on formation of the complexes DE1 and DE2, with the strongest effects observed upon removal of thymine residues at position −4 and −14 (Fig. 11c,d). In all cases, the identified negative interference effects occurred in DNA segments also protected against DNase I cleavage upon binding of the ParD2:ParE2 complex, which corroborates the specificity of the interaction (Fig. 10e).

Methylation protection experiments were performed to gather information on DNA groove-specificity of ParD2:ParE2 binding. DMS (dimethylsulphate) methylates guanine residues at the N7 position, protruding in the major groove, and adenine residues at position N3, protruding in the minor groove of the DNA helix (Siebenlist and Gilbert, 1980). Subsequent cleavage of the backbone at methylated positions by piperidine treatment essentially reveals methylated guanines, whereas the reaction rate at adenine residues is low in these conditions (Maxam and Gilbert, 1980). Binding of ParD2:ParE2 protected four guanine residues of the top strand (positions −6, −7, +4 and +5) and two of the bottom strand (positions −15 and −16) (Fig. 10c,d,e). It is worth noticing that these residues are part of inverted repeats and occur in three patches of two successive G residues, the centers of which are 9 and 10 bp apart (Fig. 10e), indicating that ParDE2:ParE2 essentially interacts with consecutive major groove segments, aligned on one face of the DNA helix.

The putative promoter of the *parDE2* operon was predicted with the software *bprom* (Solovyev and Salamov, 2011), which is a tool based on bacterial sigma 70 promoter recognition. It predicted the promoter to be at position +7 (relative to the putative start of transcription, Fig. 10e). Putative −35 (TTGCCC) and −10 (CATAAT) promoter elements separated by an ideal 17 bp stretch can be discerned slightly upstream of the transcription initiation site. The identified ParD2:ParE2 binding site completely overlaps the start of transcription (+1) and putative −10 promoter element or Pribnow box, and adjoins the −35 box of the promoter, suggesting negative autoregulation by steric exclusion of RNA polymerase binding. Therefore, our data confirm that the identified binding site corresponds to the operator region of the *parDE2* operon, which is negatively autoregulated by the ParD2:ParE2 complex in a mechanism that strongly suggests conditional cooperativity.

### ParD2 dimers in the ParD2:ParE2 complex bind DNA via the RHH motifs

The binding site of ParD2:ParE2 complex on the *parDE2* promoter extends over a 36 nt-long region, as identified by DNase I assays (see above). Interestingly, pre-modification binding interference and methylation protection experiments indicate the presence of specific protein-DNA interactions with bases in or adjacent to three imperfect palindromic repeats of sequence 5’-G[G/A]TA[C/T][C/T]-3’. SAXS measurements of ParD2:ParE2 complex bound to 31 or 21 bp DNA fragments comprising three and two palindromic repeats within the *parDE2* operator, respectively, indicate that a 6:2 T:A association binds these two DNA pieces, resulting in a protein-DNA complex of around 97 kDa with the 31 bp DNA and 91 kDa with the fragment of 21 bp (see SAXS experiments below). Therefore, model building was attempted to gain insights into the mode of DNA binding by the ParD2:ParE2 complex. By homology with the RHH family of transcriptional repressors described above, it was assumed that DNA binding occurs by insertion of the *β*-sheet formed upon ParD2 dimerization into the major groove of the DNA, and that DNA binding does not perturb the homodimeric arrangement of ParD2 molecules, as it is the case for the unliganded and DNA-bound CopG structures (Fig. 12a). The model of two ParD2 dimers bound to two 10-bp DNA regions containing two imperfect palindromic repeats of sequence 5’-G[G/A]TA[C/T][C/T]-3’ was generated based on the structure of two CopG dimers bound to its operator region (Costa et al., 2001) (Fig. 12a). First, the RHH motifs of two ParD2 dimers in the ParD2:ParE2 complex were superimposed onto the CopG dimers (Fig. 12b). Subsequently, the bases of the CopG binding site were mutated against the ParD2 site nucleotides. The sequence was then edited to match the 21 bp DNA fragment used for SAXS. To generate the longer 31 bp fragment, copies of the 21 bp were superimposed onto the already placed fragment and extended on both sides to cover the full 31 nucleotide pairs. An angle of approximately 120° was imposed across the entire fragment, as suggested by the *ab initio* models generated with the SAXS data (Fig. 13). The models were inspected for steric clashes in Chimera and energy minimization of the DNA backbone was performed when needed.

**Figure 12.**
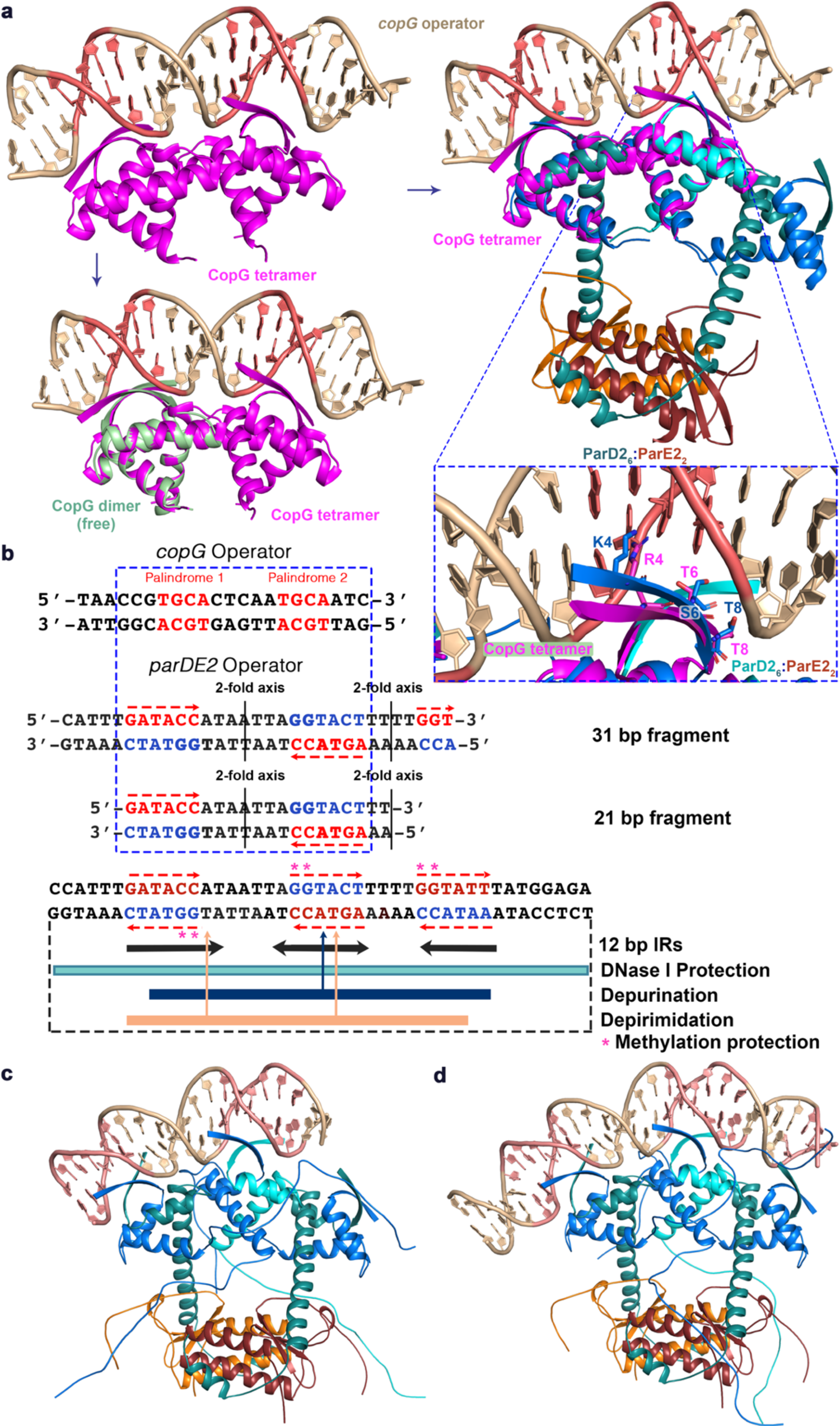
Modelling of ParD2:ParE2-Operator complex. (a) Ribbon representation of the superposition of the isolated CopG dimer onto CopG dimer in complex with a 22 bp operator DNA (PDB ID 1ea4). CopG dimerization does not change when DNA is bound. CopG dimers bind asymmetrically to a 13 bp inverted repeat comprising two short palindromic sequences. The latter are directly compared to the short imperfect palindromic repeats found in the *parDE2* operator. Fragments generated to model ParD2:ParE2 complexes with its operator DNA region as identified by DNA-interaction experiments are shown. Superposition of ParD2 homodimers in ParD2:ParE2 complex onto CopG dimers in the CopG-Operator complex. (b) Comparison between elements of symmetry in *copG* and *parDE2* operators. Ribbon representation of the final ParD2:ParE2-parDE2 Operator complexes with 21 bp-fragment (c) and 31 bp-DNA fragment (d).

**Figure 13.**
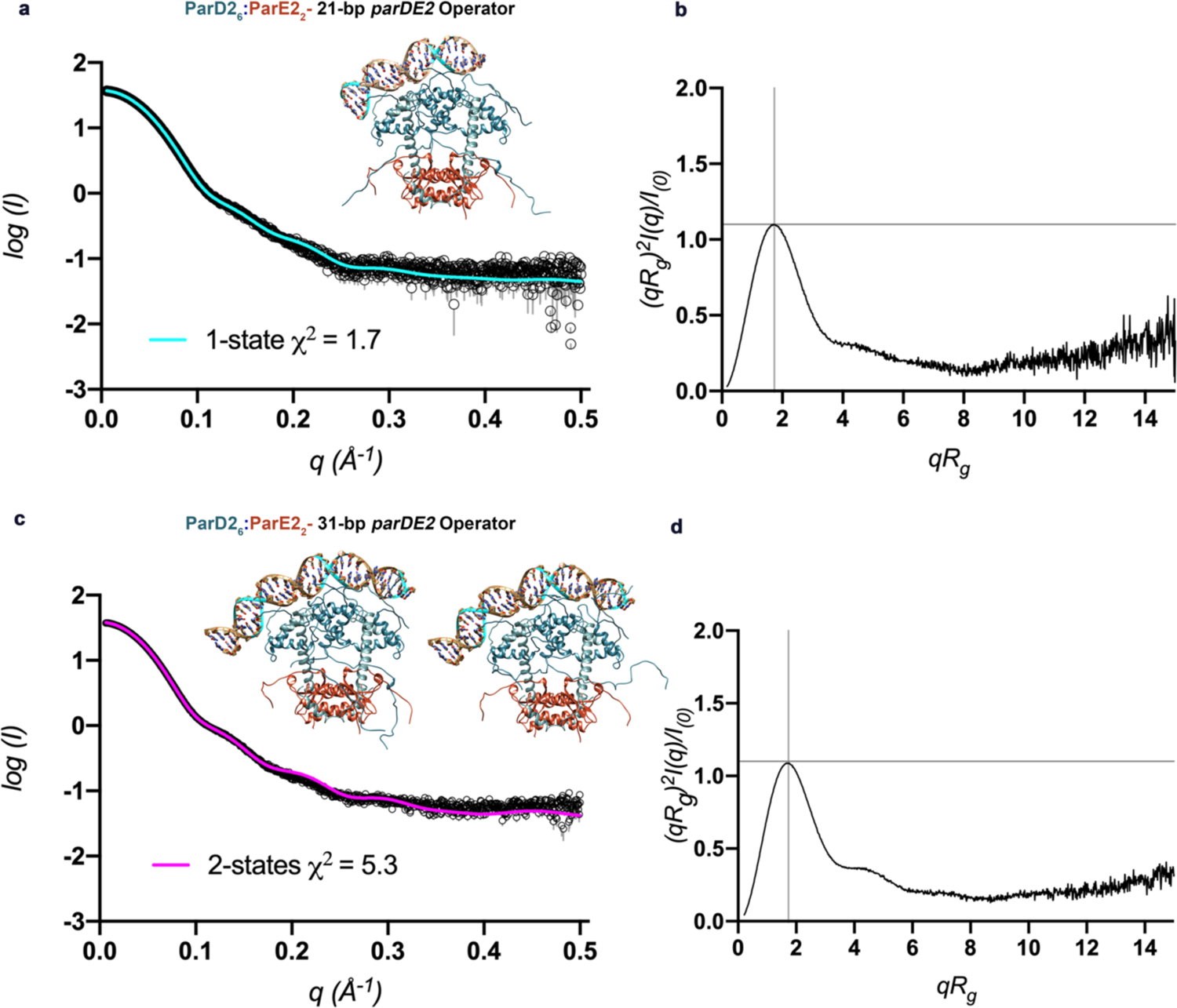
SAXS experiments validate ParD2:ParE2-DNA models. (a) Solution scattering of ParD2:ParE2 in complex with a 21 bp DNA fragment and (b) Kratky plot. The best 1-state MultiFoXS model for these complex is represented as ribbons and its theoretical SAXS profile compared to the experimental data (*χ^2^* = 1.7). (c) Experimental scattering curve for ParD2:ParE2 in complex with a 31 bp DNA fragment and Kratky plot (d). The best 2-states MultiFoXS models are represented like ribbons and their weighted theoretical profiles are compared to the experimental data (*χ^2^* = 5.3).

The *parDE2* promoter/operator region bears two 12 bp inverted repeats (IR) regions that contain three short imperfect palindromic repeats of sequence 5’-G[G/A]TA[C/T][C/T]-3’ separated by a stretch of 7 and 4 pairs of A and T nucleotides (Fig. 12c). Remarkably, our models of protein-DNA complexes reveal that these palindromic repeats are located in regions of the operator where the ParD2 homodimer *β*-sheets get inserted into consecutive major groove segments aligned on one face of the DNA helix and establish sequence specific interactions via the conserved Lys, Thr and Ser residues on the N-terminal ribbons (Fig. 12). Interestingly, as it is the case for the CopG operator interactions, two consecutive ParD2 homodimer *β*-strands contact different bases, not necessarily matching the (pseudo)-2-fold symmetry element (5’-G[G/A]TAC[C/T]-3’) of the DNA sequence. This observation that the recognition of both sides of the 12 bp inverted repeats relative to the center of the pseudo-symmetry elements is asymmetric (different protein-DNA contacts on either side), has also been made in the case of the Arc-operator complex (Raumann et al., 1994) and in the two available CopG-DNA structures (Gomis-Rüth et al., 1998; Costa et al., 2001). Additionally, the protein-DNA contacts induce a bend in the DNA which causes a compression of the minor and major grooves in the A and T stretches between the three (pseudo)-2-fold symmetry elements.

### ParD2:ParE2-DNA models are validated by SAXS

In order to validate ParD2:ParE2-DNA models, small angle X-ray scattering was measured with the protein-DNA complexes formed with the 21 and 31 bp DNA duplexes. Complexes were concentrated and injected onto a size exclusion chromatography column coupled to a capillary to which the X-rays were directed. This setup allowed for the collection of data coming from monodisperse samples of the complexes. To account for the conformational heterogeneity of the four disordered ParD2 C-termini in the comparison of the theoretical scattering of the model with the experimental SAXS data, MultiFoXS analysis was performed. The modelled-in 30 residues-long C-termini of the four ParD2 protomers in the 6:2 ParD2:ParE2 complex, together with the short C-terminal histidine-tags in both ParE2 monomers, were provided as flexible residues to MultiFoXS, while keeping the rest of the structure fixed as rigid body. 10 000 conformations were generated by MultiFoXS in each case, the theoretical SAXS profiles calculated and compared to the experimental data, and the enumeration of the best-scoring multi-state models performed. This analysis with ParD2:ParE2-21bp DNA complex yielded a best-scoring one-state model that compares to the experimental data with a *χ^2^* value of 1.70 and *R_g_* of 30.9 Å (Fig. 13a). The *R_g_* distributions across the pool of multi-state models generated by MultiFoXS overlap in the range of 29 to 32 Å, which could be correlated with the spherical shape of the ParD2:ParE2 assembly, as explained above. This result validates our ParD2:ParE2-DNA model and confirms the presence of the 6:2 TA association as the protein assembly interacting with the DNA.

Results with the 31 bp DNA complex showed a good fit to the experimental data, especially at low q values, which validates the overall shape and placement of the protein-DNA complex. In this case, the best-scoring 2 and 3-states models compare to the experimental data with *χ^2^* values of 5.40 and 5.36, respectively; with very small differences between *R_g_* values, which oscillate around 32 Å and that arise from different orientations of the disordered ParD2 C-termini in solution (Fig. 13b).

Overall, our ParD2:ParE2-DNA models indicate the ParD2-DNA mediated interactions established via ParD2 homodimer *β*-sheets, which insert into three major groove regions and contact bases in or adjacent to the (pseudo)-2-fold symmetry elements of sequence 5’-G[G/A]TA[C/T][C/T]-3’ (Fig. 13c,d).

### ParD2:ParE2 complex binds its operator with a 6:2 stoichiometry and specifically interacts with the three imperfect palindromes

In order to further validate the stoichiometry of the complex involved in DNA binding, isothermal titration calorimetry (ITC) experiments with ParD2:ParE2 complex and a 33-bp fragment containing the wild-type *parDE2* operator region were performed (Fig. 14). This fragment contains the three (pseudo)-2-fold symmetry elements (5’-G[G/A]TAC[C/T]-3’) which directly interact with three ParD2 dimers in a sequence-specific manner, as indicated in our ParD2:ParE2-DNA models described above. Binding isotherms were analyzed with the single binding site model implemented in the Microcal LLC ITC200 Origin software provided by the manufacturer. When concentrations for the ParD2:ParE2 complex are calculated in 6:2 equivalents, the *n* value of 1 obtained from the fitting to the model indicates a 1:1 stoichiometry between the ParD2:ParE2 complex and DNA fragment. The macroscopic binding constant (*K_A_*) for the binding to the wild-type (WT) operator is 4.0*10^6^ ± 4.88*10^5^ M^-1^.

**Figure 14.**
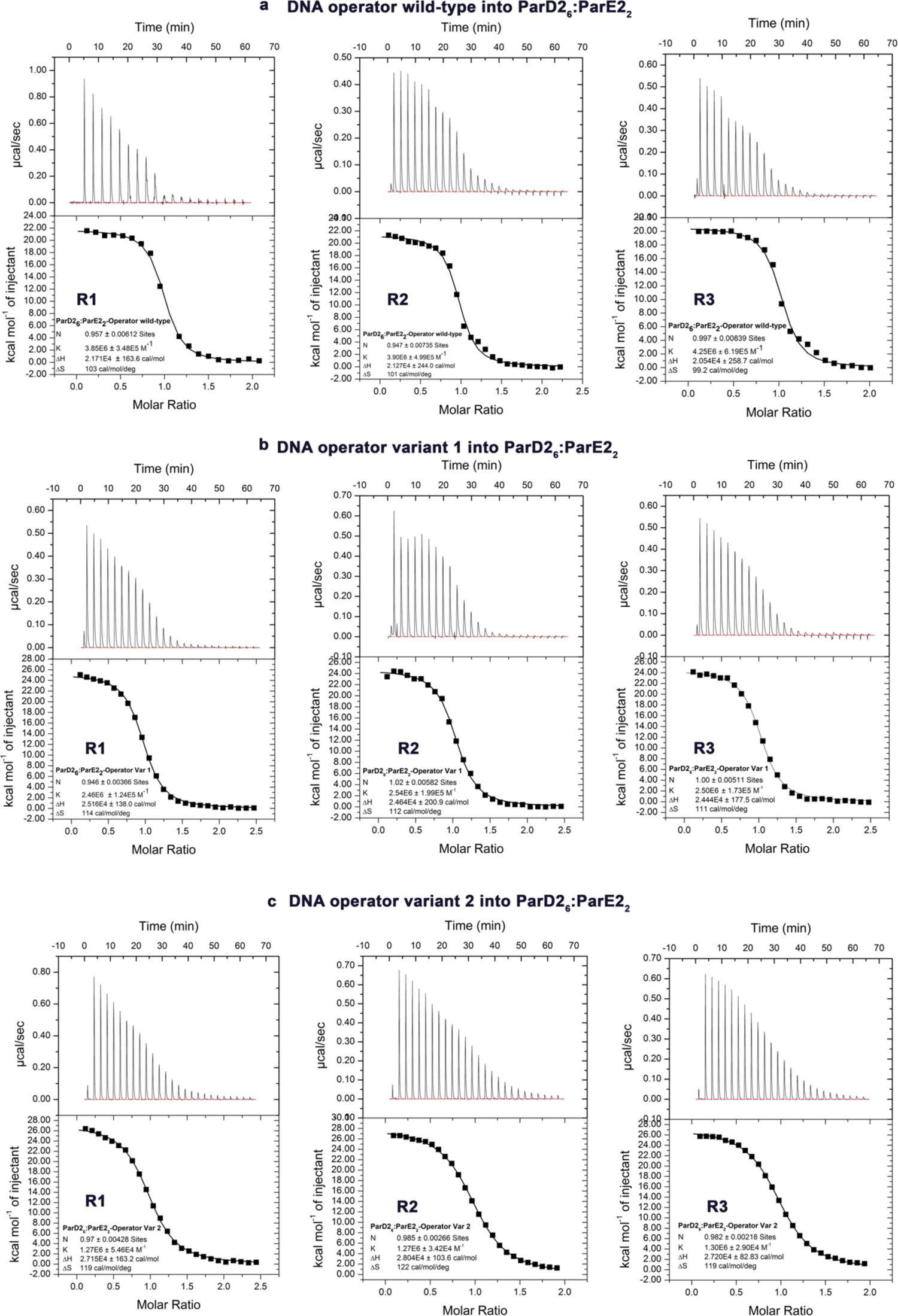
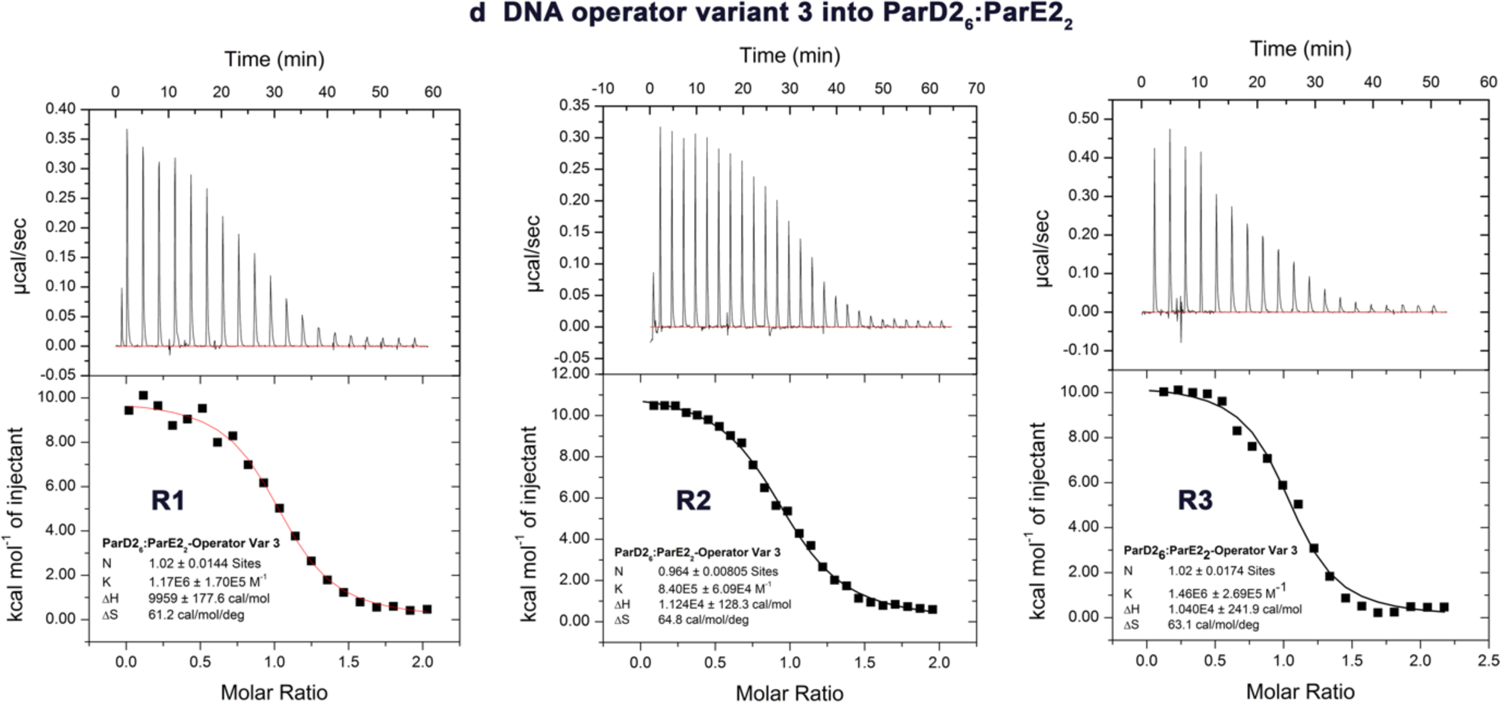
ITC titrations at 25 °C of DNA operator wild-type and variants with mutations in the short palindromic repeats into ParD2:ParE2 complex. Three replicate measurements (R1, R2, R3) of 33 bp fragment wild-type operator (a), operator variant 1 (b), operator variant 2 (c) and operator variant 3 (d).

The specificity of the protein-DNA interactions identified on the *parDE2* operator region was further assessed by ITC measurements. Four operator variants (33 bp dsDNA fragments) were designed with mutations in each of the three imperfect palindromic sequences (Table 3). Only three bases were mutated within each imperfect palindrome and ITC measurements with the *Vc*ParD2:ParE2 complex were compared to the WT fragment. The *K_A_* estimated for the binding of the *Vc*ParD2:ParE2 complex to the mutated operator variants are 4 to 1.5-fold lower than the *K_A_* estimated for the wild-type fragment. These results further confirm the specific *Vc*ParD2:ParE2 binding to the three palindromic regions. Furthermore, the stoichiometry values determined in all titrations confirm the 6:2 *Vc*ParD2:ParE2 complex as the protein assembly binding the DNA operator region. Thermodynamic parameters for each operator variant are summarized in Table 4 and represented in the graph in Fig. 15.

**Figure 15.**
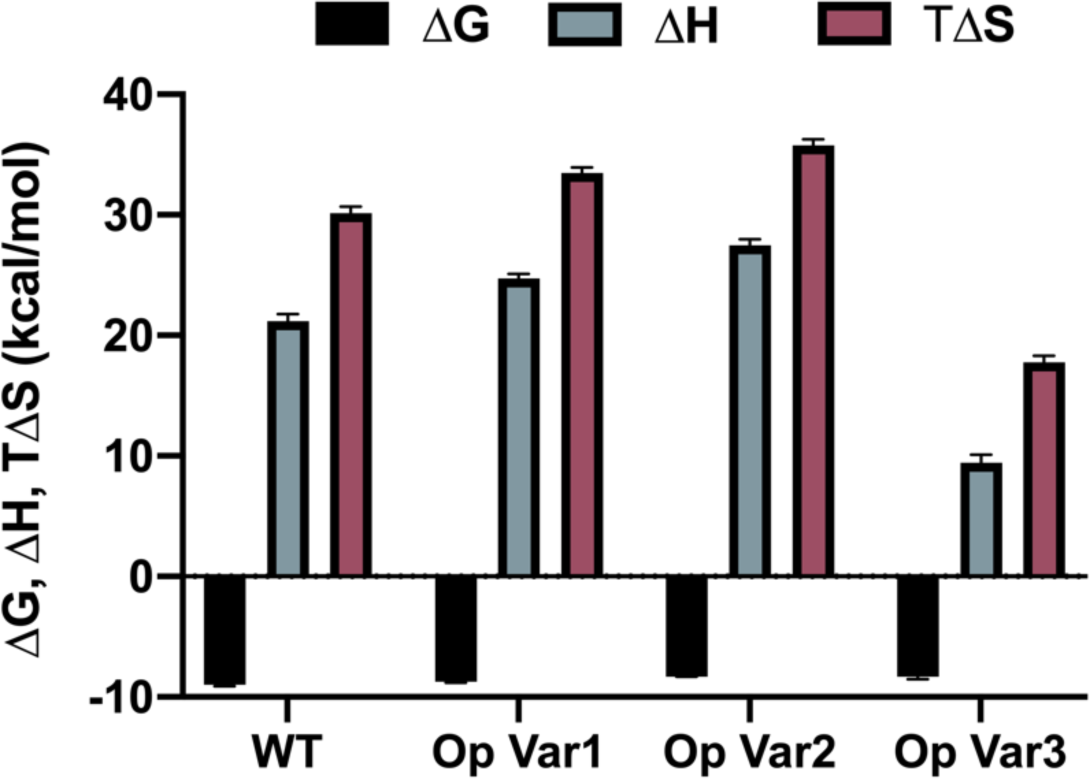
Thermodynamic parameters for the macromolecular interactions in the ParD2*6*:ParE2*2*-Operator DNA complexes (comprising wild-type (WT) and variants containing mutations in the palindromic boxes).

**Table 3.**
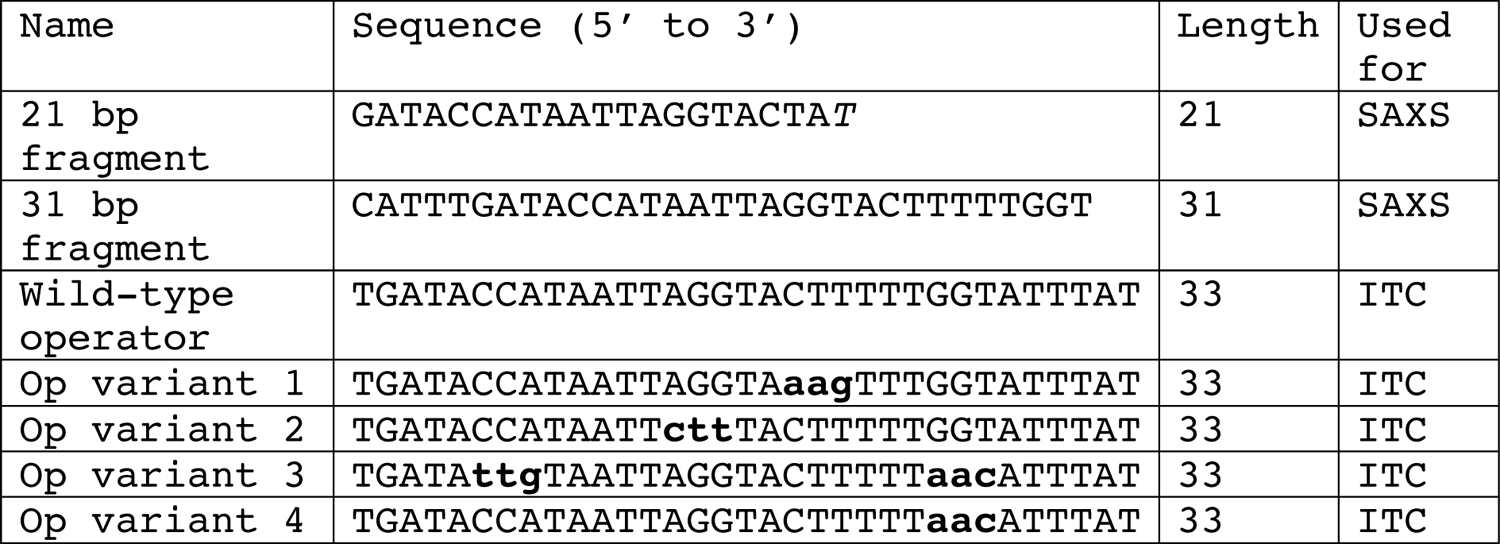
dsDNA fragments used in this study (only the top strand sequence is annotated).

**Table 4.**
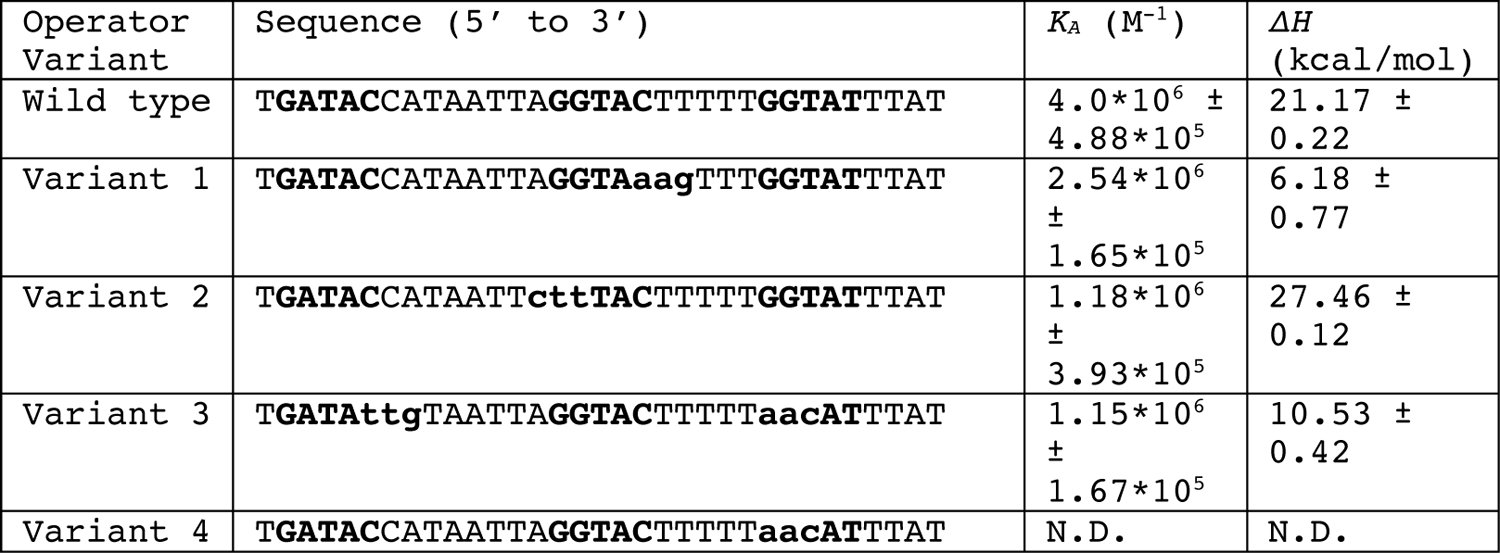
Thermodynamics of operator binding.

## Discussion

The *parDE* family is widely distributed in the genomes of bacteria and archaea, where they are also often found in multiple copies (Leplae et al., 2011). Interestingly, toxins and antitoxins from the ParDE family exhibit high interaction specificity, avoiding crosstalk between multiple paralogs in a cell. The basis of this high specificity is encoded by a small set of co-evolving residues at the toxin-antitoxin interface, allowing co-operonic pairs to exclusively interact with each other (Aakre et al., 2015). The structure of *Vibrio cholerae* ParD2:ParE2 complex presented here constitutes the third available high-resolution structure of a complex from this poorly characterized family, tallying to *C. crescentus* ParD:ParE and *M. opportunistum* ParD:ParE complexes. The structure of *E. coli* O157:H7 has been previously determined as well, however the latter deviates from the family in many aspects. *Ec*ParE does not poison DNA gyrase, nor does *Ec*ParD contain a DNA binding domain. *Ec*ParD:ParE system works as a three component TA-Regulator.

ParE toxins belong to the RelE/ParE family of toxins, and although they share the same fold, their activities and mode of antitoxin inhibition have greatly diverged during evolution. RelE toxins function as ribosome-dependent mRNases, while ParE toxins generally target DNA gyrase and interfere with replication. Structurally, there are some remarkable differences between RelE and ParE. RelE toxins contain two additional *α*-helices: short *α*-helix 3, located in the corresponding ParE loop between *α*2 and *β*2 and C-terminal *α*-helix 4, which is absent in ParE toxins. This additional C-terminal *α*-helix in RelE toxins is implicated in their toxicity. *E. coli* RelB antitoxin was shown to inhibit RelE by the displacement of *α*-helix 4 from the *β*-sheet core, which perturbs the active site configuration by sweeping away a conserved Tyr residue on *α*-helix 4 (Li et al., 2009). The structures of *E. coli* RelE in pre-and post-cleavage states bound to the 70S ribosome confirmed that isolated *E. coli* RelE does not suffer great conformational changes upon ribosome-binding and therefore resembles the active state of the toxin. The rationalization of the catalytic mechanism achieved with this study also confirmed the involvement of Tyr from *α*-helix 4 in catalysis (Neubauer et al., 2009). ParE toxins C-termini diverge from RelE, as they do not fold into an *α*-helix but remain as random coil. *Vc*ParE2 fold is highly similar to *Cc*ParE, as both of them do not contain the C-terminal *α*-helix seen in RelE. This difference might be related to the functional divergence between ParE and RelE toxins. Additionally, *Vc*ParE2, like *Cc*ParE, *Mo*ParE and *Ec*O157ParE, diverge from RelE toxins in the length of the *α*1-*α*2 hairpin. ParE toxins show a substantially longer *α*1-*α*2 hairpin compared to RelE toxins. Importantly, *Vc*ParE2, like the other ParE toxins, lacks the three crucial catalytic residues conserved in RelE homologs: Y87, R81 and R61 (Fig. 4; *E. coli* RelE numbering; Neubauer et al., 2009).

A noteworthy difference between *Vc*ParE2, *Cc*ParE, *Ec*O157ParE and the previously determined *M. opportunistum* ParE structure lies in the insertion of an 8-residues long loop between strand *β*3 and *β*4 in the latter (Pro77-Val84). *Vc*ParE2 *Cc*ParE and *Ec*O157ParE have similar *β*3-*β*4 turns, while *Mo*ParE acquired a long flexible loop that protrudes toward the solvent (Fig. 4). However, the implications of this loop in the function or specificity of these toxins is not currently understood.

The paradigm of antitoxin secondary structure induction upon toxin binding proposed and experimentally observed for many antitoxins, including RK2 ParD (Oberer et al., 2007), was questioned by Dalton and Crosson, 2010. They found *C. crescentus* ParD antitoxin to remain largely helical when not bound to *Cc*ParE toxin. Conversely, the data presented here supports the presence of intrinsic disorder in the C-terminus of *V. cholerae* ParD2 antitoxin. X-ray structures of isolated *Vc*ParD2 and in complex with *Vc*ParE2 toxin show free antitoxin chains lacking density for the last 30-residues. These same C-terminal residues then appear largely folded when bound to *Vc*ParE2 monomers in the same crystal. MS analysis confirms the presence of full-length *Vc*ParD2 antitoxin in solution, even after multiple cycles of freezing and thawing of the purified sample (data not shown), therefore ruling out the spontaneous degradation of *Vc*ParD2 C-termini. These data support the presence of diversity in the modes of antitoxin recognition and general function, more specifically that the disorder to order transition is not a universal mechanism even within this closely related antitoxins, which share high degree of structural homology, including RHH homodimerization.

Some of the most striking differences between *Vc*ParD2:ParE2 and previously determined *Cc*ParD:ParE and *Mo*ParD:ParE complexes stem from their different oligomeric assemblies. Even though all three complexes share virtually the same ParD homodimerization via their antitoxins RHH domains (homodimers superimpose with a maximum RMSD of 1.7 Å over 70 Ca), they show different assemblies in the crystals. Remarkably, both *Cc*ParD:ParE and *Mo*ParD:ParE complexes exist as saturated 1:1 stoichiometric complexes, while the stoichiometry of *Vc*ParD2:ParE2 is 3:1, with a molar excess of antitoxin. The reason for this different stoichiometries might relate to inherently different antitoxins translation rates or stability. For example, *Cc*ParD:ParE exists as an *a_2_b_2_* heterotetramer both in the crystal and in solution (Dalton and Crosson, 2010). When superimposing the RHH *Cc*ParD homodimer onto one of the *Vc*ParD2 RHH homodimers in the 6:2 assembly (Fig. 18a), clashes between toxin monomers are observed. In the saturated *Cc*ParD:ParE complex both ParE monomers are contacted by ParD chains from the same RHH homodimer. Conversely, in the 6:2 *Vc*ParD2:ParE2 assembly, ParE2 monomers are contacted by ParD2 chains from two RHH ParD2 homodimers located at opposite ends of the trimer of homodimers. In other words, the titration of the *Vc*ParD2:ParE2 complex with *Vc*ParE2 toxin, might generate a saturated heterotetramer with similar overall configuration to the one in observed in *Cc*ParD:ParE.

The differences between the two previously described complexes and *Mo*ParD:ParE are more striking. Even though the first 45 residues in both chains of *Mo*ParD RHH homodimer practically occupy the same positions as their equivalents in *Vc*ParD2 and *Cc*ParD homodimers; *α*-helix 2 in *Mo*ParD is kinked around R47, causing it to deviate from a straight helix path (Fig. 16b). Approximately a 10-15° angle between the trajectories of *Vc*ParD2 *α*2 and *Mo*ParD *α*2 is generated by the kink in *Mo*ParD *α*2. This causes a displacement of *Mo*ParE monomers, which no longer contact each other in the *Mo*ParD:ParE heterotetramer (Fig. 16b). It could be compared to the “opening” of the *Cc*ParD:ParE heterotetramer by pulling ParE toxins apart in opposite directions. Moreover, this much more open heterotetramer is not present in solution as such (according to analytical exclusion chromatography assays), but rather as a dimer of tetramers (Aakre et al., 2015). In that assembly two heterotetramers are associated via ParE interactions, in a sort of complementary “Lego”, thereby decreasing the solvent accessible area of the whole assembly. This hetero-octamer is also the most stable assembly output by the PISA server analysis of this structure (PDB ID 5ceg), and as expected, the buried area of 35260 Å^2^ upon the formation of the hetero-octamer is almost 3-fold higher than the buried area of the heterotetramer. The different assemblies observed in these three complexes indicate that, despite the high homology in the fold of ParD and ParE proteins across species, they can adopt different oligomerization states and topologies, which could correlate with intrinsic differences in the transcriptional repression mechanisms of the ParD:ParE family.

**Figure 16.**
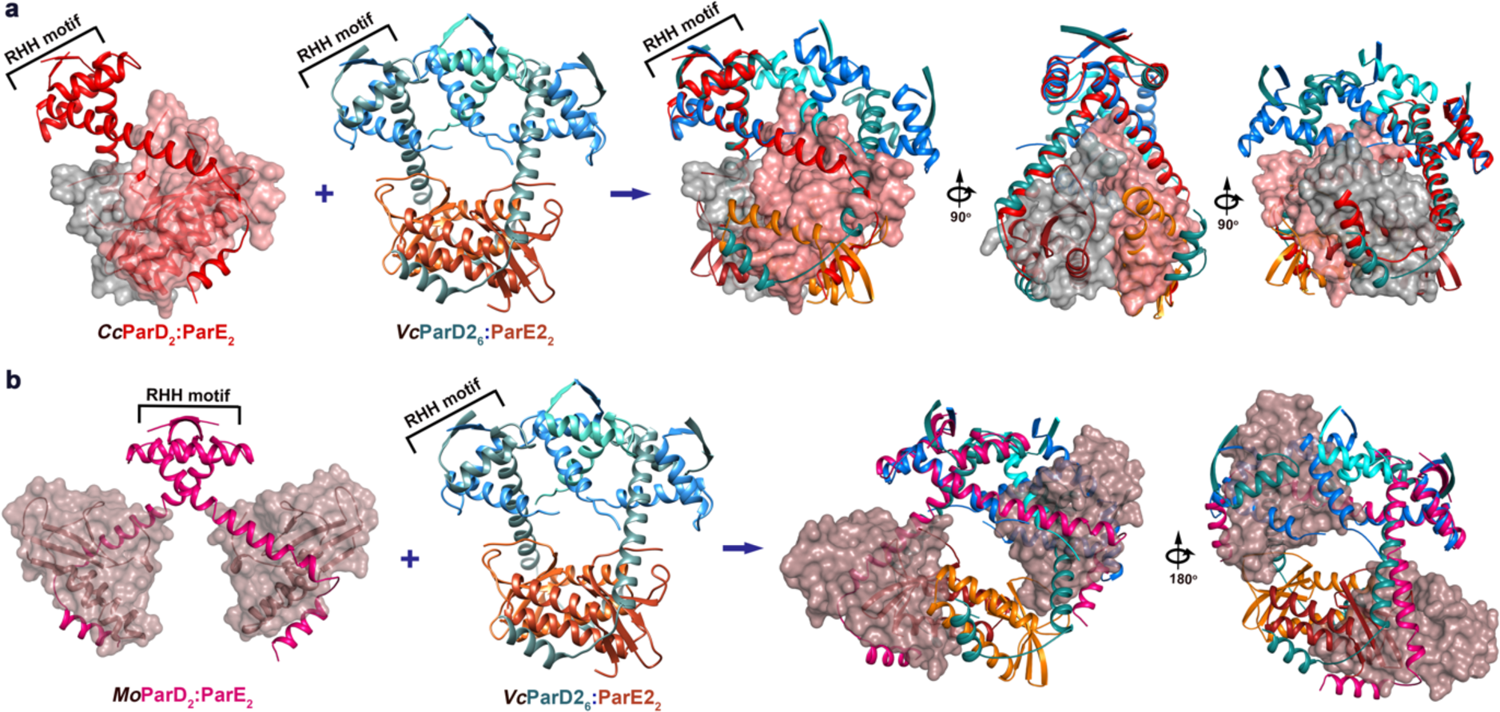
Comparison between *V. cholerae* ParD2:ParE2, *C. crescentus* ParD:ParE (PDB ID 3kxe) and *M. opportunistum* ParD:ParE (PDB ID 5ceg) complex assemblies. Ribbon representation of (a) *C. crescentus Cc*ParD:ParE and (b) *M. opportunistum Mo*ParD:ParE heterotetramers and their superimposition onto *V. cholerae Vc*ParD2:ParE2 hetero-octamer via RHH homodimerizing domains on ParD antitoxins. ParE toxins from *C. crescentus* and *M. opportunistum* are represented as surface.

The specific DNA sequence contacted by the *Vc*ParD2:ParE2 complex was experimentally determined by DNase I footprinting experiments, methylation protection assays and pre-modification binding interference assays. The same 151-bp labelled DNA fragment was used across different assays and DNA strand-specific interactions were identified by using top and bottom-labelled DNA. These experiments permitted the identification of a 36 nt-long DNA region which is specifically protected by *Vc*ParD2:ParE2 complex from DNase I. Importantly, all other techniques identified protein-DNA contacts comprised within this same region, which confirmed the specificity and accuracy of the experimentally determined *parDE2* operator sequence. Additionally, methylation protection experiments indicate the specific recognition of the guanines in the three imperfect palindromes of sequence 5’-G[G/A]TA[C/T][C/T]-3’, which further supports the *Vc*ParD2:ParE2-DNA binding mode proposed in this work. Moreover, pre-modification binding interference experiments additionally showed protein-specific interactions with several bases within or adjacent to the three imperfect palindromes identified in the *parDE2* operator region. Particularly clear are interaction with thymines −4 and −14 and adenine −5 on the bottom strand, which are within the central palindrome and the palindrome upstream the latter, respectively. Moreover, when analyzing the DNA sequence, the −10 and −35 promoter elements were clearly identified upstream the *parD2* translation start. The identified operator region completely overlaps with the −10 box and adjoins the −35 element, thereby supporting the mechanism by which *Vc*ParD2:ParE2 binding the operator region would abrogate RNA polymerase binding the promoter and thereby repress transcription. ITC experiments further confirmed the specificity of the binding via the three imperfect palindromes and the stoichiometry of the binding supports the 6:2 *Vc*ParD2:ParE2 assembly binding the DNA operator region.

Structurally, *Vc*ParD2 is classified into the MetJ/Arc/CopG family of transcriptional repressors, along with its RK2 homolog ParD. The archetype short transcriptional repressor CopG shows striking similarities to *Vc*ParD, both in protein fold and homodimer topology. Furthermore, the crystallographic contacts made by CopG also show a second dimerization interface between adjacent molecules, analogous to what is observed in *Vc*ParD2. These interactions in the crystal render a super helical structure formed by 12 CopG monomers per turn and the RHH DNA-binding regions all directed toward the outside. CopG super-helical assembly is analogous to the disk-like shape adopted by isolated *Vc*ParD2 (RHH motifs located on the disk’s mantle) and the 120° angle formed by three adjacent N-terminal ParD2 dimers in the ParD2:ParE2 complex structure. The average distance between two adjacent *β*-ribbons (two dimers) formed upon ParD2 and CopG dimerization via the RHH motifs is approximately 37 Å in both models, analogous to the inter-RHH distance in the *Vc*ParD2:ParE2 crystallographic assembly. Interestingly, this dimer of homodimers in the CopG structure does not show significant deviations between the unliganded CopG crystal and the DNA-bound CopG structure, supporting that DNA binding does not induce significant changes in the protein structure. These observations indicate that the formation of the second inter-homodimer interface might be conserved within this family of repressors, especially in those proteins with conserved Gly-mediated turns. Furthermore, the formation of the higher-order repressor assemblies could work as a fine-tuned mechanism to control repression levels without a big genetic load for the cell, as proposed by del Solar et al., 2002.

Based on the homology between *Vc*ParD2 and CopG, homology models for the binding of *Vc*ParD2:ParE2 to DNA fragments of different lengths encoding the *parDE2* operator region were generated and validated by SAXS. The available crystallographic structure of a CopG tetramer bound to a 22bp DNA fragment (PDB ID 1ea4) was used as starting model. There are some differences between the *copG* operator and *parDE2* operator region. For example, *copG* operator comprises two 4 bp perfect palindromes of sequence 5’-TGCA-3’ which are separated by 5 bp. *parDE2* operator region on the other hand, is comprised of three 6 bp imperfect palindromes of sequence 5’-G[G/A]TA[C/T][C/T]-3’ which are separated by 7 and 4 bp regions. However, the crystallographic structure of CopG-22 bp DNA shows asymmetry in protein-DNA specific interactions. For example, RHH residue R4, which stablishes the most base-specific interactions in the CopG-DNA complex, contacts the DNA asymmetrically. Two R4 from the same dimer contact different bases on the DNA, e.g., R4 on CopG chain G contacts the two CG bases on the left of the 5’-TGCA-3’ palindrome on the coding strand, while it also contacts the palindromic adenine base on the bottom strand. However, R4 from the other chain of the same homodimer (chain F), makes the most contacts with the palindromic bases (three last bases of the 5’-TGCA-3’ on the top strand and two bases within the palindrome on the bottom strand). Analogously, the adjacent homodimer makes contacts with the bases flanking the palindromic regions toward the distal end of the DNA fragment. This observation supports the idea that the palindromic regions are important for specificity, however, the binding is not limited to the palindrome bases and therefore not necessarily symmetric. Another evidence in support of this mode of DNA recognition and binding was presented as a model for the interaction of CopG-DNA. Costas et al., 2001 observed in DNase I and hydroxyl radical footprinting experiments (with plasmid linear fragments containing the inverted repeats in a centered position), the successive protection of 4-5 nt regions separated by 6-7 nt unprotected zones, even at relatively low protein concentrations. This finding supports the cooperative binding of CopG homodimers on the same DNA face, which are able to nucleate along the DNA as protein concentration increases.

When analyzing the symmetry in the *parDE2* operator region, the different distance between inverted repeats stands out as a noticeable feature. However, unlike the *copG* operator, all the bases located in the region between the three imperfect palindromes in the *parDE2* operator exclusively comprise A and T bases. A-T pairs are easily compressed due to the absence of bulky exocyclic groups in the minor groove of the helix. Furthermore, when there are at least 4 consecutive A’s or T’s the DNA may adopt the B’ form stabilized by propeller twisting and characterized by a narrower minor groove and intrinsic bending (McConnell and Beveridge, 2001). The linker between palindromes (boxes) 2 and 3 has such a stretch of 4 Ts in a row. Therefore, the AT-rich regions between the (pseudo)-2-fold symmetry elements allow for a higher “plasticity” in the DNA conformation of this region.

We tested that hypothesis by measuring SAXS of *Vc*ParD2:ParE2-DNA complexes. Fragments of 21 bp and 31 bp comprising two and three imperfect palindromes, respectively, were used for SAXS measurements. The first evidence toward the binding of the 6:2 *Vc*ParD2:ParE2 assembly to the dsDNA fragment stemmed from the estimated molecular masses of the protein-DNA complexes, which agreed with the expected values for the binding of the hetero-octameric assembly. Furthermore, the homology models for the two fragments generated from the CopG-operator structure were refined to account for the conformational heterogeneity of the 4 *Vc*ParD2 disordered tails in the hetero-octamer. The calculated theoretical scattering curves for the refined models show high agreement to the measured SAXS curves for the two protein-DNA complexes. This further supports the mode of binding of *Vc*ParD2:ParE2 to its DNA operator region via the three RHH motifs on the surface of the antitoxin.

The basis for the conditional cooperativity observed in the EMSAs could stem from the *Vc*ParD2 inter-homodimer contacts. This suggests a highly cooperative binding that is further stabilized by *Vc*ParE2 at high antitoxin:toxin ratios. The relevance of the 3:1 antitoxin:toxin ratio for the optimal DNA-binding becomes clear when looking at the 6:2 *Vc*ParD2:ParE2 assembly. This assembly allows for the preferential stabilization of the *Vc*ParD2 trimer of homodimers, which are pre-positioned within the complex for optimal binding to the three imperfect palindromes of the *parDE2* operator. This model explains our DNase I footprinting results, which show the same protected region on the DNA at all tested *Vc*ParD2:ParE2 concentrations. Moreover, the SEC-MALS results showed one single peak with the same molecular mass corresponding to the 6:2 *Vc*ParD2:ParE2 assembly, in all measured concentration in the range of 0.25-10 mg ml^-1^, indicating that the complex preferentially adopts this conformation. These results taken together suggest that even at the comparatively low protein concentrations used for DNase I footprinting (the highest protein concentration there being 0.15 mg ml^-1^), 6:2 *Vc*ParD2:ParE2 complex binds and thereby protects the 36-nt operator DNA region with high affinity and specificity. Interestingly, weak protection is observed in DNase I footprinting at 0.1 mmol l^-1^ (0.0075 mg ml^-1^) ParD2:ParE2 concentration, although hyperreactivity effects are marked (12a). This could indicate binding of excess free ParD2 homodimers rather than the 6:2 complex, which were detected in the co-purified ParD2:ParE2 sample by native MS (Fig. 1). Binding of ParD2 homodimers at different sites (palindromes or boxes) of the operator with comparable affinity could cause only a fraction of the molecules to be protected at a specific site, therefore protection by DNase I would be masked (which agrees with the absence of protected region when isolated ParD2 antitoxin is used in DNase I assays). It cannot be ruled out that the reason for the lack of visible protected region from DNase I when using isolated ParD2 antitoxin could also be fast DNA dissociation rates. Subsequently, when antitoxin:toxin ratios decrease, by addition of ParE2 toxin, the 6:2 ParD2:ParE2 assembly becomes destabilized. Free toxin monomers bind the flexible ParD2 tails protruding on both sides of the assembly which causes the folding upon binding of ParD2 C-termini, thereby causing steric clashes with previously bound toxin molecules. This titration process drives the disassembly of the 6:2 ParD2:ParE2 complex via the rupture of ParD2 inter-homodimer contacts, generating for example, a ParD2:ParE2 hetero-tetramer topologically similar to the *C. crescentus* complex, as discussed above (Fig. 17; see diagram on Fig. 18). If such a complex is superimposed onto one of the two CopG dimers in complex with its 22 bp operator fragment, it becomes clear that two ParD:ParE heterotetramers could not consecutively bind two sites on the operator due to steric hindrance (Fig. 17). Therefore, cooperative binding mediated by ParD2 inter-homodimer contacts would become abrogated. The binding affinity of a ParD2 homodimer in the heterotetramer for a single DNA site might be too low for efficient DNA binding and repression, as suggested by our toxin-titration EMSAs (Fig. 9b).

**Figure 17.**
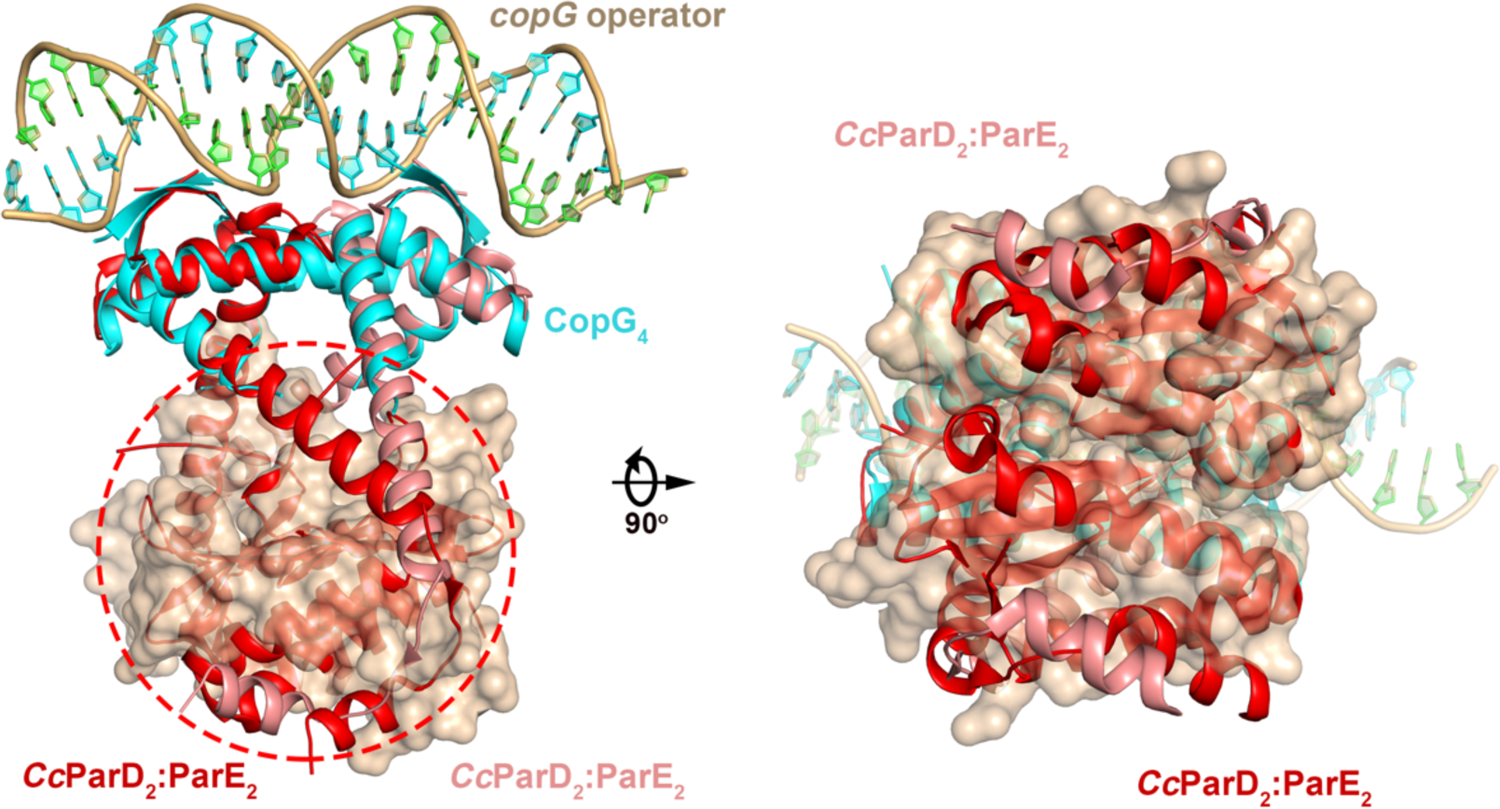
Consecutive ParD2:ParE2 heterotetramers cannot simultaneously bind *parDE2* operator region due to major steric clashes. Ribbon representation of CopG-operator (CopG is color cyan) (PDB ID 1ea4) with two *C. crescentus* ParD:ParE heterotetramers superposed onto the CopG homotetramer. One *Cc*ParD:ParE complex is colored red and the other one pink. ParE toxins from the pink *Cc*ParD:ParE complex are represented as surface. The area with major clashes between complexes is encircled.

These results taken together permit the assignment of the different species observed on the EMSAs with ParD2:ParE2 complex and isolated ParD2 antitoxin. DE1 and DE2 could correspond to the binding of one and two ParD2 homodimers on different operator boxes, respectively (Fig. 8, Fig. 9). Strikingly, these two bands show the same mobility as the D1 and D2 on the EMSA performed with isolated ParD2 antitoxin. At low ParD2:ParE2 complex concentrations, the equilibrium is shifted toward the dissociation of the 6:2 assembly into free ParD2 antitoxins, which coexist with the complex in solution as observed by native MS analysis of the copurified complex (Fig. 1). When total protein concentration increases, the equilibrium shifts to the highly stable 6:2 ParD2:ParE2 assembly, which readily binds DNA as three consecutive ParD2 homodimers are pre-positioned for binding the three consecutive imperfect palindromes. This generates a super-shifted complex with high molecular weight: DE3. This observation lays the basis for the conditional cooperativity observed in the EMSAs with increasing toxin:antitoxin ratios. When 0.5 μmol l^-1^ of ParD2:ParE2 starting concentration is used (equivalent to 3 μM ParD2), DE3 complex is formed (Fig. 8c, Fig. 9a). As toxin:antitoxin ratios increase, DNA binding transitions to DE2 species. This could potentially arise from the released ParD2 homodimers or ParD2:ParE2 hetero-tetramer binding the operator, as the 6:2 ParD2:ParE2 assemble dissociates (Fig. 9a, diagram on Fig. 17). Interestingly, when ParD2:ParE2 starting concentration is halved (0.25 μmol l^-1^), the visible band corresponds to DE2 due to the dilution of the complex. As the toxin:antitoxin ratio increases, ParD2 is titrated into the ParD2:ParE2 heterotetramer, with the concomitant appearance of DE1 (ParD2 dimer binding DNA), culminating in total DNA release when all ParD2:ParE2 exists as heterotetramers in solution (Fig. 9, Fig. 17).

**Figure 18.**
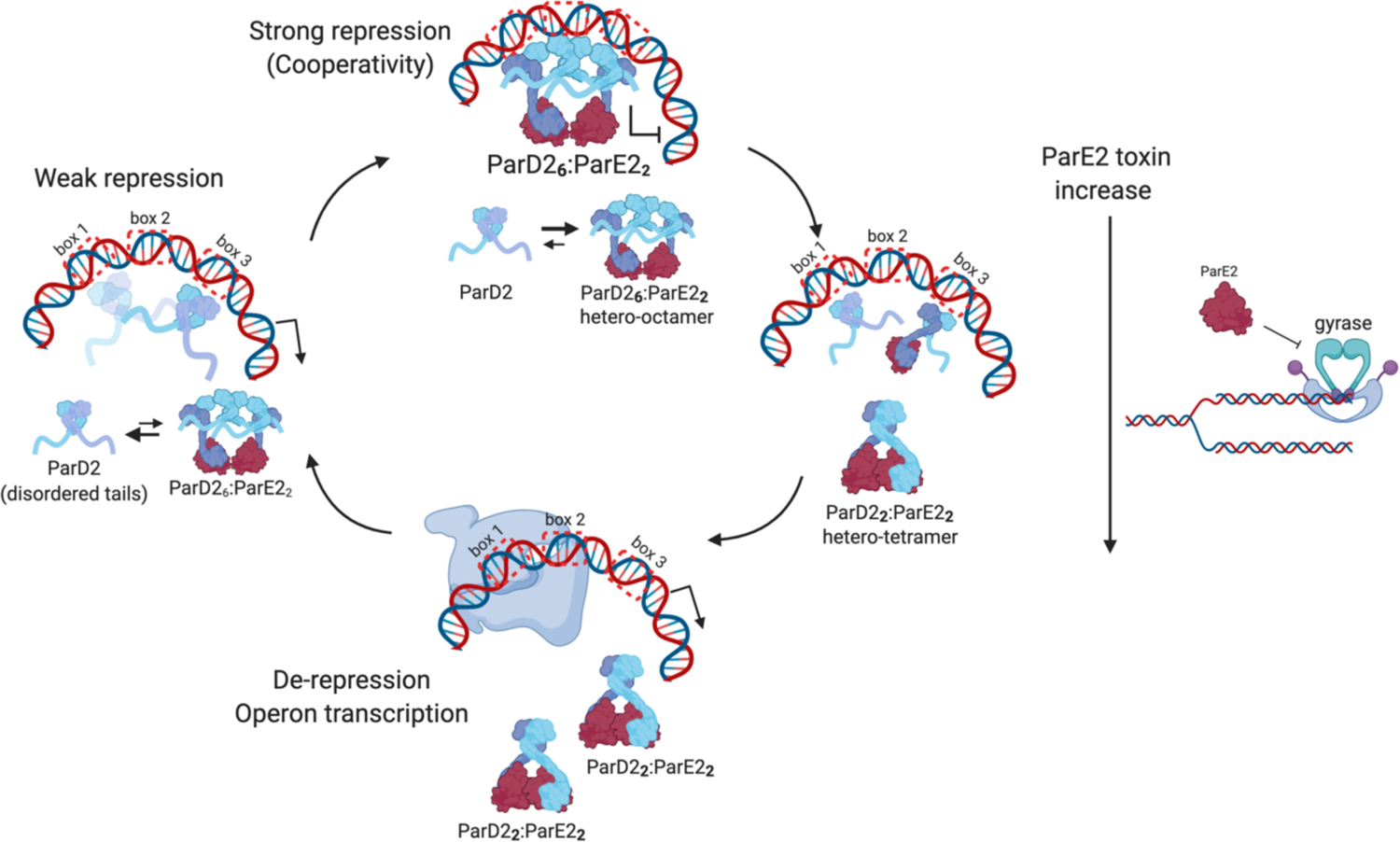
Regulation of the *V. cholerae parDE2* operon. ParD2 antitoxin (shown in shades of blue) exists as RHH homodimers with disordered C-termini in solution. At low toxin:antitoxin ratios or absence of toxin, ParD2 homodimers bind operator boxes (imperfect palindromic repeats) with high dissociation rates or low affinity. Binding of ParE2 induces the structuring of ParD2 C-termini and the preferential stabilization of the 6:2 ParD2:ParE2 assembly, which pre-orients three antitoxin homodimers in the right orientation for DNA binding via simultaneous contacts with the three operator boxes. This complex causes the strongest operator binding and thereby the highest repression of transcription initiation. When toxin:antitoxin ratios increase, ParE2 toxins bind ParD2 C-termini and break the 6:2 assembly. ParD2 antitoxin is liberated and able to weakly bind the operator, therefore restoring some level of transcription. The saturated 1:1 TA heterotetramer is not able to consecutively bind the three operator boxes due to steric hindrance. When all TA complex exists as the saturated 1:1 TA heterotetramer, operator binding is abrogated, and transcription rates increase to replenish the antitoxin pool.

## Methods

### Cloning, protein expression and purification of ParD2:ParE2-His

The coding region of the *parDE2* operon from *V. cholerae* biovar El Tor strain N16961 (NCBI NC_002506.1) fused to a histidine-tag coding strip placed C-terminal to *parE2* was synthesized by Genscript (https://www.genscript.com/). This gene was subsequently cloned into the NcoI-NheI sites of a pET28a plasmid and verified by DNA sequencing. *E. coli* BL21 (DE3) competent cells (William Studier et al., 1990) were transformed with the expression plasmid using the CaCl_2_ method (Hanahan et al., 1991). Transformed colonies were selected on LB plates containing kanamycin (50 mg ml^-1^) and grown overnight at 310 K. LB medium supplemented with kanamycin was inoculated with one isolated colony and grown overnight at 310 K with aeration. A 100-fold dilution of the former preculture was used to start 1 l LB cultures containing 50 mg ml^-1^ kanamycin. Cells were grown at 310 K with aeration until OD_600_ reached 0.6. The temperature was then decreased to 293 K and protein expression was induced by adding 0.5 mM isopropyl *β*-D-1-thiogalactopyranoside (IPTG). After overnight incubation at 293 K with aeration, cells were harvested by centrifugation at 277 K for 15 min at 6500 g (5000 rev min^-1^ in a JLA-8.1000 rotor) and subsequently resuspended in lysis buffer (20 mM Tris-HCl pH 8, 500 mM NaCl, 2 mM *β*-mercaptoethanol) supplemented with a protease-inhibitor cocktail (cOmplete, Sigma-Aldrich/Merck) and stored at 193 K. The cell suspensions were incubated with DNase I (50 μg ml^-1^) and MgCl_2_ (2 mM) for 10 min at room temperature before extraction with a continuous-flow cell disruptor (Constant Systems) at 277 K and a pressure between 17-18 KPSI. Cell debris was removed by centrifugation at 41 700g (18 000 rev min^-1^ in a JA-20 rotor) for 50 min at 277 K. The cleared extract was filtered (0.45 μm) before loading onto a nickel-nitrilotriacetic acid affinity column (connected to an AKTAexplorer FPLC system, GE Healthcare) previously equilibrated with lysis buffer. The loaded fraction was subsequently washed with the same lysis buffer to remove non-specific contaminants. ParD2-ParE2 complex was eluted using a step imidazole gradient (10, 25, 50, 250 and 500 mM) in 20 mM Tris–HCl pH 8, 500 mM NaCl, 2 mM *β*-mercaptoethanol over 5 column volumes per imidazole concentration. The progress of the purification was analysed via SDS-PAGE (Laemmli, 1970) and Western blot (Towbin et al., 1979), with a mouse anti-histidine antibody (BioRad, AbD Serotec). The ParD2:ParE2 containing fraction was subsequently applied onto a pre-equilibrated Superdex 200 (16/60) column (GE Healthcare) in SEC buffer (20 mM Tris–HCl pH 8, 150 mM NaCl, 1 mM tris(2-carboxyethyl)phosphine) and run at 1.0 ml min^-1^. The fractions containing pure ParD2:ParE2 complex were pooled, flash-frozen and stored at 193 K.

### Denaturant-induced dissociation of the ParD2:ParE2-His complex

The ParD2-ParE2 on-column denaturant-induced dissociation was adapted and optimized from the protocol for *E. coli* O157 PaaA2-ParE2 TA system described by (Sterckx et al., 2015). Briefly, the pure ParD2-ParE2 complex in SEC buffer was loaded onto a nickel–nitrilotriacetic acid affinity column (connected to an AKTAexplorer FPLC system, GE Healthcare) previously equilibrated in the same SEC buffer. The loaded ParD2-ParE2-His complex was then washed with buffer A2 (20 mM Tris-HCl pH 8, 1 M NaCl, 10 % ethylene glycol) for 5 column volumes (CV). ParD2 was then eluted with GnHCl over a two-step gradient: 50 and 100 % of buffer A3 (20 mM Tris-HCl pH 7, 500 mM NaCl, 5 M GnHCl) for 10 CV each. The fractions containing ParD2 were pooled and dialyzed overnight at 277 K in SEC buffer (supplemented with 2 mM *β*-mercaptopethanol instead of TECP). These fractions were concentrated and applied onto a Superdex 200 (16/60) column in SEC buffer as a final polishing step. The fractions containing ParD2 were pooled, flashed-frozen and stored at 193 K.

The Ni-NTA bound and unfolded ParE2-His was refolded on column by the sequential application of 10 CV of each of the following buffers: low salt buffer A4 (25 mM Tris pH 8, 25 mM NaCl, 2 mM *β*-mercaptoethanol, 5 % glycerol), higher salt buffer A5 (25 mM Tris pH 8, 250 mM NaCl, 2 mM *β*-mercaptoethanol, 1 % glycerol) and A6 (25 mM Tris pH 8, 250 mM NaCl, 2 mM *β*-mercaptoethanol). The refolded toxin was then eluted by a step-gradient of imidazole with buffer A7 (20 mM Tris pH 8, 500 mM NaCl, 500 mM imidazole, 2 mM *β*-mercaptoethanol). The final polishing step consisted of a gel filtration on Superdex 75 (16/60) in SEC buffer. The fractions containing ParE2 were pooled, flashed-frozen and stored at 193 K.

### Generation, expression and purification of Nanobodies against ParD2:ParE2 complex

ParD2:ParE2 specific nanobodies were generated as described before (Pardon et al., 2014). In brief, a total of 0.5 mg ml^-1^ of pure ParD2-ParE2 complex was used for the immunization of a llama (*Lama glama*). Four days after the final boost, blood was taken to isolate peripheral blood lymphocytes. RNA was purified from these lymphocytes and reversed transcribed by PCR to obtain cDNA. The resulting library was cloned into the phage display vector pMESy4 bearing a C-terminal hexa-His tag and a Glu-Pro-Glu-Ala-tag (also called EPEA-tag or CaptureSelect C-tag). Phage display selections against ParD2:ParE2 were performed using two different approaches. First, ParD2:ParE2 was solid phase coated in NaHCO_3_ pH 8.2 and selections were done in 20 mM Tris pH 8, 150 mM NaCl, 1 mM TCEP either in the absence or presence of soluble ParD2. All clones were screened on ParD2:ParE2 solid phase coated. Screening was performed in 20 mM Tris pH8, 150 mM NaCl, with 1 mM TCEP when Nanobodies were present. For the detection of the presence of specific Nanobodies, the “EPEA”-tag at the end of the Nanobody was used. The CaptureSelect Biotin anti-C-tag Conjugate (LifeTechnologies #7103252100; 1/4000) was mixed with Streptavidin Alkaline Phosphatase (Promega #V5591; 1/1000 dilution) for 10 minutes before use. Signal was detected using 4-Nitrophenyl phosphate disodium salt hexahydrate (2 mg ml^-1^). In total, 92 clones were screened in ELISA, 58 clones were positive. The 58 clones were sequence analysed and 34 families were found. If several variants of a family are found, more than one is included. A Nanobody family is defined as a group of Nanobody amino acid sequences with high similarity in the CDR3 sequence. By default, we consider nanobodies belonging to the same family as binding to the same target epitope.

Plasmids were transformed to *E. coli* WK6 cells and checked by DNA sequencing after transformation. Nanobodies were expressed and purified according to steps 70-73 in the protocol described (Pardon et al., 2014). Briefly, pre-cultures were prepared by inoculating a single transformed *E. coli* WK6 colony into a 50-ml sterile Falcon tube containing 10 mL of LB supplemented with 100 μg ml^-1^ ampicillin, 2% (wt/vol) glucose and 1 mM MgCl_2_. Precultures were grown overnight at 310 K with aeration. 1 l bottles of TB media supplemented with 100 μg ml^-1^ ampicillin and 1 mM MgCl_2_ were inoculated with the 10 ml precultures. The bottles were incubated at 310 K with aeration until OD_600_ reached 0.7. Then, protein expression was induced by adding 1 mM IPTG and the cultures were incubated at 301 K with aeration for 20 h. Cells were collected by centrifugation for 10 min at 5000 rev min^-1^ in a JLA-8.1000 rotor at 277 K. The pellet from each 1 l bottle was resuspended in 15 ml ice-cold TES buffer and incubated for 1 hour in an orbital shaking platform at 277 K. 30 ml of TES/4 buffer were added to the resuspended pellets and further incubated for 45 min on ice while shaking. The suspension was centrifuged for 40 min at 5000 rev min^-1^ in a JLA-8.1000 rotor at 277 K and the supernatant was recovered as the periplasmic extract.

For purification of the nanobodies, a first IMAC step was performed followed by size exclusion chromatography. Ni-NTA beads were resuspended three times in 50 ml of wash buffer (50 mM phosphate buffer, 1 M NaCl pH 7) at RT. The periplasmic extract was added to the washed Ni-NTA beads and incubated for 1 hour at RT on a shaking platform. This mix was added to a PD 10 column and subsequently washed with 10 ml of wash buffer (pH 7), followed by 20 ml of wash buffer at pH 6. The elution of nanobody was done by adding 10 ml of 50 mM acetate buffer, 1 M NaCl pH 4.5 to the column and immediately neutralizing the elution in fractionation tubes containing 2 ml of 1 M Tris-HCl pH 7.4. The eluted fractions were concentrated up to 2 ml and applied onto a Superdex 75 16/60 column equilibrated with 20 mM Tris pH 8, 150 mM NaCl. The purity of the eluted fractions was assessed by SDS-PAGE and Western blot using a mouse anti-histidine antibody (BioRad, AbD Serotec).

### Native Mass spectrometry

Spectra were recorded on a traveling wave Q-TOF instrument (Synapt G2, Waters, Manchester, UK) tuned for transmission of large, native protein assemblies. Samples were introduced into the gas phase at a concentration of 1.8 μM and 7 μM for ParD2:ParE2 complex and ParD2 antitoxin, respectively, in 150 mM ammonium acetate pH 7.8 using nano-electrospray ionization with in-house prepared gold-coated borosilicate glass capillaries. Critical voltages throughout the instrument were a sampling cone voltage of 25 V and a trap collision energy of 10 V (unless mentioned otherwise), with pressures throughout the instrument of 5.7 and 2.25 E-2 mbar for the source and trap collision cell regions. Spectra were externally calibrated using a 10 mg/mL solution of cesium iodide. Analyses of the acquired spectra were performed using Masslynx version 4.1 (Waters, Manchester, UK). Native MS spectra were smoothed (extent depending on size of the complexes) and additionally centered for calculating molecular weights.

### Crystallization, X-Ray data collection and processing

ParD2:ParE2-His in 20 mM Tris–HCl pH 8, 150 mM NaCl, 1 mM tris(2-carboxyethyl)phosphine was concentrated to 12 mg ml^-1^. To make super-complexes with nanobodies, concentrated ParD2:ParE2 was mixed with a molar excess of nanobody and incubated for 30 min at RT. The mix was subsequently injected onto a Superdex 200 16/60 equilibrated in SEC buffer. Fractions corresponding to ParD2:ParE2:Nb super-complexes were pooled and concentrated. SDS-PAGE was performed to check for the formation of super-complexes.

Crystallization conditions were screened using a Mosquito HTS robot from TTP Labtech (http://ttplabtech.com/) using 0.1 μl protein solution and 0.1 μl of reservoir solution equilibrated against 100 μl of reservoir solution in sitting drop configuration and later on incubated at 292 K. Crystals grew after two years in 0.1 M KCl, 0.1 M Na-HEPES pH 7, 15 % PEG 5000 MME. For data collection, crystals were flash-cooled in liquid nitrogen. Data were collected on PROXIMA-2A at the SOLEIL synchrotron facility, Gif-sur-Yvette, France. All data were indexed, integrated and scaled with XDS (Kabsch, 2010). Data quality and twinning were analyzed with *phenix.xtriage* (Liebschner et al., 2019) and POINTLESS (Evans, 2006). Analysis of solvent contents was performed using the CCP4 program MATTHEWS_COEF (Kantardjieff & Rupp, 2003).

### Structure determination

The structure of ParD2:ParE2 complex was determined by molecular replacement in Phaser-MR. Several truncations of the *C. crescentus* ParD:ParE model (PDB ID 3kxe) were employed as search models. However, only a single molecule of *C. crescentus* ParE could be placed in the AU with LLG and TFZ values of 34 and 8.4, respectively. In spite of the low LLG value, this partial solution was refined in *phenix.refine* using rigid body refinement and manually refined in Coot to mutate the amino acid sequence to that of *Vc*ParE2. This refined VcParE2 model was defined as known partial solution for another Phaser-MR search with the N-terminal *Vc*ParD2 dimer. One copy of the dimer was placed, but the AU was still not complete. A final Phaser-MR search defining the previous partial solution with the full-length *C. crescentus* ParD monomer, placed a third ParD molecule in the AU. Further iterative cycles of *phenix.refine* including NCS, TLS, and secondary structure restraints were performed combined with manual model building in Coot.

The structure of ParD2:ParE2:Nb80 supercomplex was determined by molecular replacement with Phaser-MR using the previously determined *Vc*ParD2:ParE2 structure and a homology model of Nb80. Iterative cycles of *phenix.refine* and manual model building in *Coot* were done as described for the crystal structure of ParD2:ParE2. Data collection and refinement statistics for all structures are given in Table 1.

### Protein_DNA interaction studies

A 350 bp-long DNA fragment comprising the first 67 bp of the *parD2* ORF and extending further upstream in the control region of the *parDE2* operon from *V. cholerae* biovar El Tor strain N16961 (NCBI NC_002506.1) was synthesized by PCR-based gene assembly (Stemmer et al., 1995) upon optimization of the oligonucleotides in the DNAWorks platform (Hoover and Lubkowski, 2002). The assembled 350 bp fragment was used as the template for PCR amplification of a 151 bp-long fragment comprising the putative operator region upstream of *parD2* ORF with the oligonucleotides forward (Fw) 5’-tgaggcgtttgttatgcgc and reverse (Rv) 5’-tttgtatttggcttgtaataaagccat as primers, of which one was [5’-^32^P] single-end labelled with (γ ^32^P)-ATP (Perkin Elmer, 3000 Ci.mmol^-1^) and T4 polynucleotide kinase (Thermofisher) as described (Nguyen Le Minh et al., 2018). Labelled PCR fragments were purified by gel electrophoresis on 6% polyacrylamide. For EMSAs, ParD2:ParE2, ParD2, ParE2 and reconstituted ParD2:ParE2 complex were mixed with labelled DNA (10 000 – 15 000 cpm) in ParD2:ParE2 buffer (20 mM Tris pH 8, 150 mM NaCl, 1 mM TCEP) in a total volume of 20 μl and incubated at 20°C for 30 min. All binding assays were performed in the presence of an excess non-specific, non-labelled competitor DNA (25 μg ml^−1^ herring sperm DNA), unless otherwise indicated. After incubation, 3 μl loading buffer (25 % ficoll, 0.1% xylenexyanol and 0.1% bromophenol) were added to each sample. Separation was performed on 6% polyacrylamide gels run in TBE buffer at 130 V for approximately 3 hours.

*DNase I footprinting* experiments with purified ParD2:ParE2 complex were performed by the method of (Galas and Schmitz), 1978, in ParD2:ParE2 buffer (see above) as described by (Charlier and Boyen’) Charlier et al., 1992. In brief, Fw* and Rv* [5’-^32^P]-labelled DNA fragments (150 000 cpm) were incubated with 0.1, 0.5, 1 and 2 μmol l^-1^ ParD2:ParE2 respectively, as described above. DNase I was then added to a final concentration of 0.025 units ml^-1^ and the reaction was stopped after 5 mins by the addition of 12.5 μl of stop mix (3 M ammonium acetate, 0.25 M EDTA), 30 μl H_2_O and 3 μl of yeast tRNA (5 mg ml^-1^). The DNA was subsequently precipitated with ethanol and the reaction products analysed by 10% (Fw*-labelled) or 6% (Rv*-labelled) denaturing polyacrylamide gel electrophoresis.

Sparingly modified [5’-^32^P] single-end labelled DNA (on average one modification per DNA molecule) for use in premodification binding interference (missing contact) assays (Brunelle and Schleif, 1987) were generated as described (Maxam and Gilbert, 1980). Premodified Fw* (top) and Rv* (bottom) labelled-DNA was incubated with 0.1, 0.5 and 2 μmol l^-1^ of ParD2:ParE2 complex, respectively, as described above for EMSAs, and analysed by native gel electrophoresis. The resolved free and bound DNA forms corresponding to different complexes were recovered from gel, cleaved at modified positions by piperidine-treatment (Maxam and Gilbert, 1980) and the reaction products subsequently analysed by denaturing gel electrophoresis as described above.

*Methylation protection* experiments were performed as described (Wang) (Wang et al., 1998). 0.1, 0.25, 0.5, 1 and 2 μmol l^-1^ of ParD2:ParE2 complex was mixed with single end Fw* (top) and Rv* (bottom) labelled, respectively, and incubated as described above. Limited methylation was performed by the addition of 1 μl DMS for one min at 20°C. The reaction was stopped with 50 μl of stop solution (1.5 M sodium acetate (pH 7.0), 1.0 M 2-mercaptoethanol) and 15 μg of yeast tRNA. Piperidine-induced strand scission was performed before analysis by denaturing gel electrophoresis. Reference ladders were generated by chemical sequencing of the 151 bp labelled fragments (Maxam and Gilbert, 1980) and all gels were autoradiographed to display the bands.

### Preparation of double-stranded oligonucleotides

The wild type *parDE* operator region and its variants were generated from single-stranded DNA oligonucleotides (Sigma). Equimolar amounts of complementary oligonucleotides were mixed in water and incubated at 95 °C for 5 mins. Subsequently, samples were allowed to slowly cool down to room temperature for approximately 1 hour. The annealing was verified by analytical size exclusion chromatography.

### Isothermal Titration Calorimetry

ITC titrations were performed in an iTC200 calorimeter (GE Healthcare) following overnight dialysis of ParD2:ParE2 complex, ParD2 antitoxin and all DNA variants in 20 mM Tris pH 8, 150 mM NaCl, 1 mM TCEP. Before experiments, samples were degassed for 5 mins and all titrations were performed at 25 °C. ParD2_6_:ParE2_2_ complex and ParD2_6_ antitoxin at 24,3 μM and 27,5 μM respectively, were added into the 200 ul cell. 2 ul injections of WT *parDE* operator at 240,5 μM were performed.

Raw data were integrated and corrected for the buffer dilution heat effects using the MicroCal Origin software to obtain the enthalpy change per mole of added ligand as a function of DNA/protein ratio. Calorimetric isotherms were analyzed with the single binding site model (concentration values in 6:2 equivalents of ParD2:ParE2 complex) using the Microcal LLC ITC200 Origin software provided by the manufacturer.

### Modeling of the ParD2:ParE2-DNA complex

Modelling of ParD2:ParE2-DNA was performed with the structure of *V. cholerae* ParD2:ParE2 complex and the available structure of the transcriptional repressor CopG from *Streptococcus agalactiae* plasmid PMV158, in complex with its 22 bp-DNA binding site (PDB ID 1ea4). The backbone of the ParD2 dimer was superimposed onto the template RHH dimer of CopG (resid 1-40) in Pymol. The two palindromic sequences (5’-TGCA-3’) separated by 5 bp in the CopG operator DNA were mutated to the two imperfect palindromes in the *parDE2* operator (5’-GTA[C/T]-3’) separated by 6 bp. The remaining CopG operator DNA sequence was mutated to the corresponding bases in the *parDE2* operator in Chimera. Subsequently, a copy of this DNA fragment was used to extend the model to a 31 bp fragment, maintaining an angle of approximately 120° on the DNA fragment. Phosphodiester bonds to connect adjacent fragments were created in Pymol and the sequence edited to the corresponding *parDE2* operator fragment of 31 bp comprising the three imperfect palindromic repeats identified by DNase I protection assays and used for SAXS measurements. The DNA structure was regularized by energy minimization in Chimera and combined with an all-atoms model of ParD2:ParE2 complex. Hydrogen atoms were added to both ParD2:ParE2-DNA models, and later used for comparison to the SAXS data in MultiFoXS (Schneidman-Duhovny et al., 2016).

### Small angle X-ray scattering

SAXS data were collected at beamline BM29 (ESRF) in HPLC mode. Shodex KW402.5-4F column was used to collect data for ParD2:ParE2. Superdex 200 Increase column (GE Healthcare) was used to collect data for ParD2:ParE2-DNA complexes and DNA samples. Protein samples were prepared as described above for crystallization, concentrated and briefly spun down before loading on the SEC column. 25 μl of 18 mg ml^-1^ ParD2:ParE2 were injected onto the Shodex column.

Double-stranded DNA fragments were generated as described above. 100 μM ParD2:ParE2 (6:2) complex were incubated with 50 μM of 31 bp-DNA or 21 bp-DNA operator variants in 500 μl of final volume. Protein-DNA complexes were incubated at 25 °C for 15 mins before injection onto a Superdex 200 Increase gel filtration column (GE Healthcare). The peak corresponding to DNA-protein complex was collected and concentrated. 75 μl of the concentrated ParD2:ParE2-DNA peak and 100 μM DNA samples were injected onto the Superdex 200 Increase 200 connected to the capillary to which the X-rays were directed.

All samples were measured in 20 mM Tris, pH 8, 150 mM NaCl, 1 mM TCEP at 19 °C. Constant column flows of 0.2 ml min^-1^ and 0.75 ml min^-1^ were used for the Shodex KW402.5-4F and the Superdex 200 Increase columns, respectively. The final scattering curve (after buffer subtraction) was generated for each sample after a range of scattering curves around the peak (with equivalent *R_g_* values) was normalized and averaged. The *R_g_* values were derived from the Guinier approximation at small *q* values while the *I_o_* parameter was estimated by extrapolation to *q* = 0 using the ATSAS suite (Franke et al., 2017). Molecular weights were determined by the Bayesian estimation implemented in Primus (ATSAS suite).

### SAXS data analysis

Computation of the SAXS profile of the atomic resolution models and its comparison to the experimental data was performed by FoXS webserver (Schneidman-Duhovny et al., 2016) (https://modbase.compbio.ucsf.edu/foxs/), which optimizes the excluded volume (*c_1_*) and hydration layer density (*c_2_*) to improve the fit of a model to the experimental SAXS data.

The MutiFoXS tool of this webserver was used to address conformational heterogeneity in solution by considering multiple states contributing to the observed SAXS profile.

All-atoms input models for ParD2:ParE2 complex and ParD2:ParE2-DNA complexes generated as described above were used for the calculations. 10 000 conformations were generated by sampling the space of the φ and ψ main chain dihedral angles of selected flexible residues with a Rapidly exploring Random Trees (RRTs) algorithm. Disordered ParD2 C-termini residues were defined as flexible while keeping the rest of the protein or protein-DNA complex as rigid body. Subsequently, the computation of the SAXS profile for each sampled conformation was performed followed by the enumeration of the best-scoring multi-state models using the multi-state scoring function implemented in MultiFoXS (Schneidman-Duhovny et al., 2016).

